# How the Avidity of Polymerase Binding to the -35/-10 Promoter Sites Affects Gene Expression

**DOI:** 10.1101/597989

**Authors:** Tal Einav, Rob Phillips

**Author notes:** Corresponding author; 1200 E. California Blvd, MC 103-33, Pasadena, CA 91125 Corresponding author; 1200 E. California Blvd, MC 128-95, Pasadena, CA 91125.

## Abstract

Although the key promoter elements necessary to drive transcription in *Escherichia coli* have long been understood, we still cannot predict the behavior of arbitrary novel promoters, hampering our ability to characterize the myriad of sequenced regulatory architectures as well as to design novel synthetic circuits. This work builds on a beautiful recent experiment by Urtecho *et al.* who measured the gene expression of over 10,000 promoters spanning all possible combinations of a small set of regulatory elements. Using this data, we demonstrate that a central claim in energy matrix models of gene expression – that each promoter element contributes independently and additively to gene expression – contradicts experimental measurements. We propose that a key missing ingredient from such models is the avidity between the -35 and -10 RNA polymerase binding sites and develop what we call a *refined energy matrix* model that incorporates this effect. We show that this the refined energy matrix model can characterize the full suite of gene expression data and explore several applications of this framework, namely, how multivalent binding at the -35 and -10 sites can buffer RNAP kinetics against mutations and how promoters that bind overly tightly to RNA polymerase can inhibit gene expression. The success of our approach suggests that avidity represents a key physical principle governing the interaction of RNA polymerase to its promoter.

**Significance Statement:** Cellular behavior is ultimately governed by the genetic program encoded in its DNA and through the arsenal of molecular machines that actively transcribe its genes, yet we lack the ability to predict how an arbitrary DNA sequence will perform. To that end, we analyze the performance of over 10,000 regulatory sequences and develop a model that can predict the behavior of any sequence based on its composition. By considering promoters that only vary by one or two elements, we can characterize how different components interact, providing fundamental insights into the mechanisms of transcription.

## Introduction

Promoters modulate the complex interplay of RNA polymerase (RNAP) and transcription factor binding that ultimately regulates gene expression. While our knowledge of the molecular players that mediate these processes constantly improves, more than half of all promoters in *Escherichia coli* still have no annotated transcription factors in RegulonDB (1) and our ability to design novel promoters that elicit a target level of gene expression remains limited.

As a step towards taming the vastness and complexity of sequence space, the recent development of massively parallel reporter assays has enabled entire libraries of promoter mutants to be simultaneously measured (2–4). Given this surge in experimental prowess, the time is ripe to reexamine how well our models of gene expression can extrapolate the response of a general promoter.

A common approach to quantifying gene expression, called the *energy matrix model*, assumes that every promoter element contributes additively and independently to the total RNAP (or transcription factor) binding energy (3). This model treats all base pairs on an equal footing and does not incorporate mechanistic details of RNAP-promoter interactions such as its strong binding primarily at the -35 and -10 binding motifs (shown in Fig. 1A). A newer method recently took the opposite viewpoint, designing an RNAP energy matrix that only includes the -35 element, -10 element, and the length of the spacer separating them (5), neglecting the sequence composition of the spacer or the surrounding promoter region.

**Figure 1.**
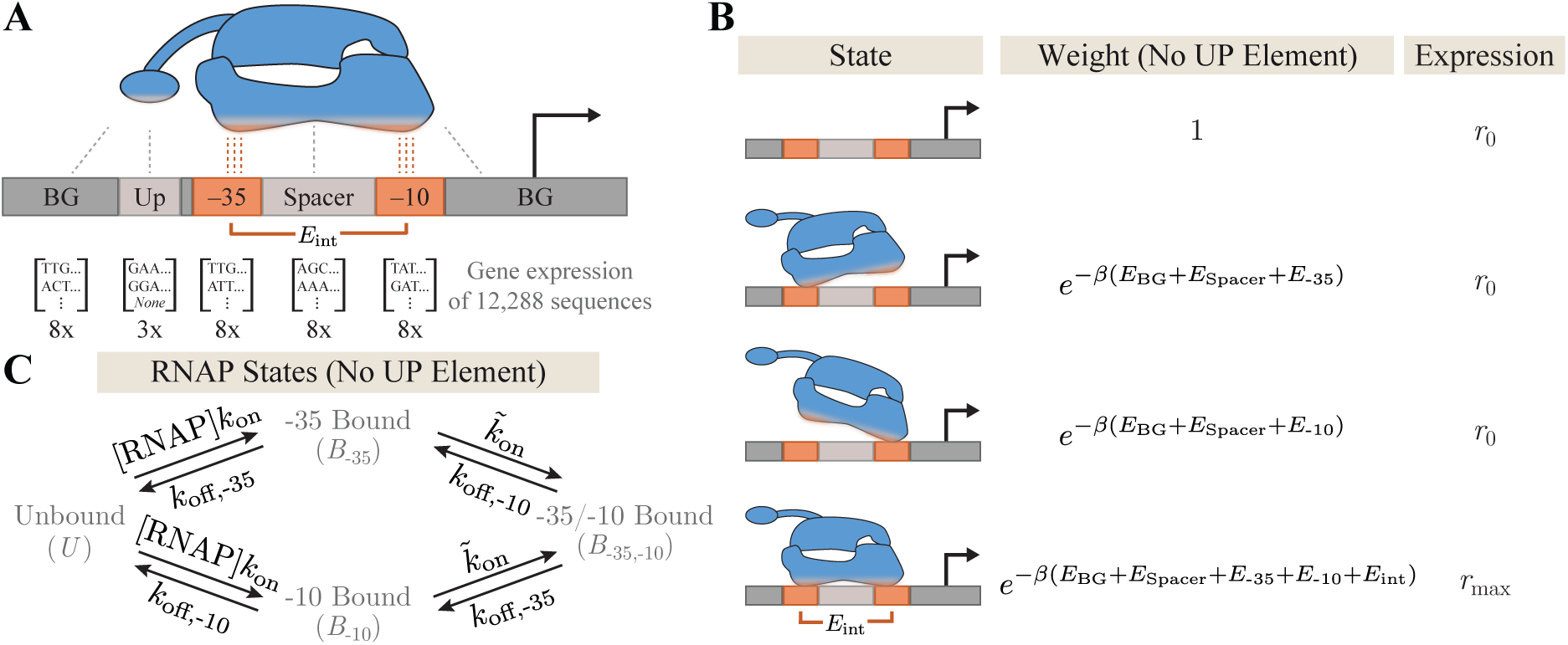
The bivalent nature of RNAP-promoter binding. (A) Gene expression was measured for RNAP promoters comprising any combination of -35, -10, spacer, UP, and background (BG) elements. (B) When no UP element is present, RNAP makes contact with the promoter at the -35 and -10 sites giving rise to gene expression *r*_0_ when unbound or partially bound and *r*_max_ when fully bound. (C) Having two binding sites alters the dynamics of RNAP binding. *k*_on_ represents the on-rate from unbound to partially bound RNAP and 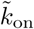 the analogous rate from partially to fully bound RNAP, while *k*_off,*j*_ denotes the unbinding rate from site *j*.

Although these methods have been successfully used to identify important regulatory elements in unannotated promoters (6) and predict evolutionary trajectories (5), it is clear that there is more to the story. Even in the simple case of the highly-studied *lac* promoter, such energy matrices show systematic deviations from measured levels of gene expressions, indicating that some fundamental component of transcriptional regulation is still missing (7).

We propose that one failure of current models lies in their tacit assumption that every promoter element contributes independently to the RNAP binding energy. By naturally relaxing this assumption to include the important effects of avidity, we can push beyond the traditional energy matrix analysis in several key ways including: (*i*) We can identify which promoter elements contribute independently or cooperatively without recourse to fitting, thereby building an unbiased mechanistic model for systems that bind at multiple sites. (*ii*) Applying this approach to RNAP-promoter binding reveals that the -35 and -10 motifs bind cooperatively, a feature that we attribute to avidity. Moreover, we show that models that instead assume the -35 and -10 elements contribute additively and independently sharply contradict the available data. (*iii*) We show that the remaining promoter elements (the spacer, UP, and background shown in Fig. 1A) do contribute independently and additively to the RNAP binding energy and formulate the corresponding model for transcriptional regulation that we call a *refined energy matrix model*. (*iv*) We use this model to explore how the interactions between the -35 and -10 elements can buffer RNAP kinetics against mutations. (*v*) We analyze a surprising feature of the data where overly-tight RNAP-promoter binding can lead to decreased gene expression. (*vi*) We validate our model by analyzing the gene expression of over 10,000 promoters in *E. coli* recently published by Urtecho *et al.* (8) and demonstrate that our framework markedly improves upon the traditional energy matrix analysis.

While this work focuses on RNAP-promoter binding, its implications extend to general regulatory architectures involving multiple tight-binding elements including transcriptional activators that make contact with RNAP (CRP in the *lac* promoter (9)), transcription factors that oligomerize (as recently identified for the *xylE* promoter (6)), and transcription factors that bind to multiple sites on the promoter (DNA looping mediated by the Lac repressor (10)). More generally, this approach of categorizing which binding elements behave independently (without resorting to fitting) can be applied to multivalent interactions in other biological contexts including novel materials, scaffolds, and synthetic switches (12, 13).

## Results

### The -35 and -10 Binding Sites give rise to Gene Expression that Defies Characterization as Independent and Additive Components

Decades of research have shed light upon the exquisite biomolecular details involved in bacterial transcriptional regulation via the family of RNAP *σ* factors (14). In this work, we restrict our attention to the *σ*^70^ holoenzyme (8), the most active form under standard *E. coli* growth condition, whose interaction with a promoter includes direct contact with the -35 and -10 motifs (two hexamers centered roughly 10 and 35 bases upstream of the transcription start site), a spacer region separating these two motifs, an UP element just upstream of the -35 motif that anchors the C-terminal domain (*α*CTD) of RNAP, and the background promoter sequence surrounding these elements.

Urtecho *et al.* constructed a library of promoters composed of every combination of eight -35 motifs, eight -10 motifs, eight spacers, eight backgrounds (BG), and three UP elements (Fig. 1A) (8). Each sequence was integrated at the same locus within the *E. coli* genome and transcription was quantified via DNA barcoding and RNA sequencing. One of the three UP elements considered was the absence of an UP binding motif, and this case will serve as the starting point for our analysis.

The traditional energy matrix approach used by Urtecho *et al.* posits that every base pair of the promoter will contribute additively and independently to the RNAP binding energy (8), which by appropriately grouping base pairs is equivalent to stating that the free energy of RNAP binding will be the sum of its contributions from the background, spacer, -35, and -10 elements (see Appendix A). Hence, the gene expression (GE) is given by the Boltzmann factor

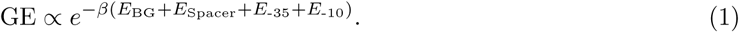

Note that all *E*_*j*_s represent free energies (with an energetic and entropic component); to see the explicit dependence on RNAP copy number, refer to Appendix A. Fitting the 32 free energies (one for each background, spacer, -35, and -10 element) and the constant of proportionality in Eq. 1 enables us to predict the expression of 8 × 8 × 8 × 8 = 4,096 promoters.

Fig. 2A demonstrates that Eq. 1 leads to a poor characterization of these promoters (*R*^2^ = 0.57, parameter values listed in Appendix B), suggesting that critical features of gene expression are missing from this model. One possible resolution is to assumes that the level of gene expression saturates for very strong promoters at *r*_max_ and for very weak promoters at *r*_0_ (caused by background noise or serendipitous near-consensus sequences (5)), namely,

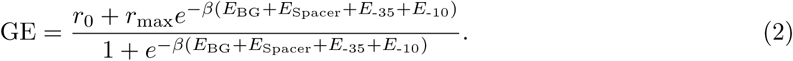

**Figure 2.**
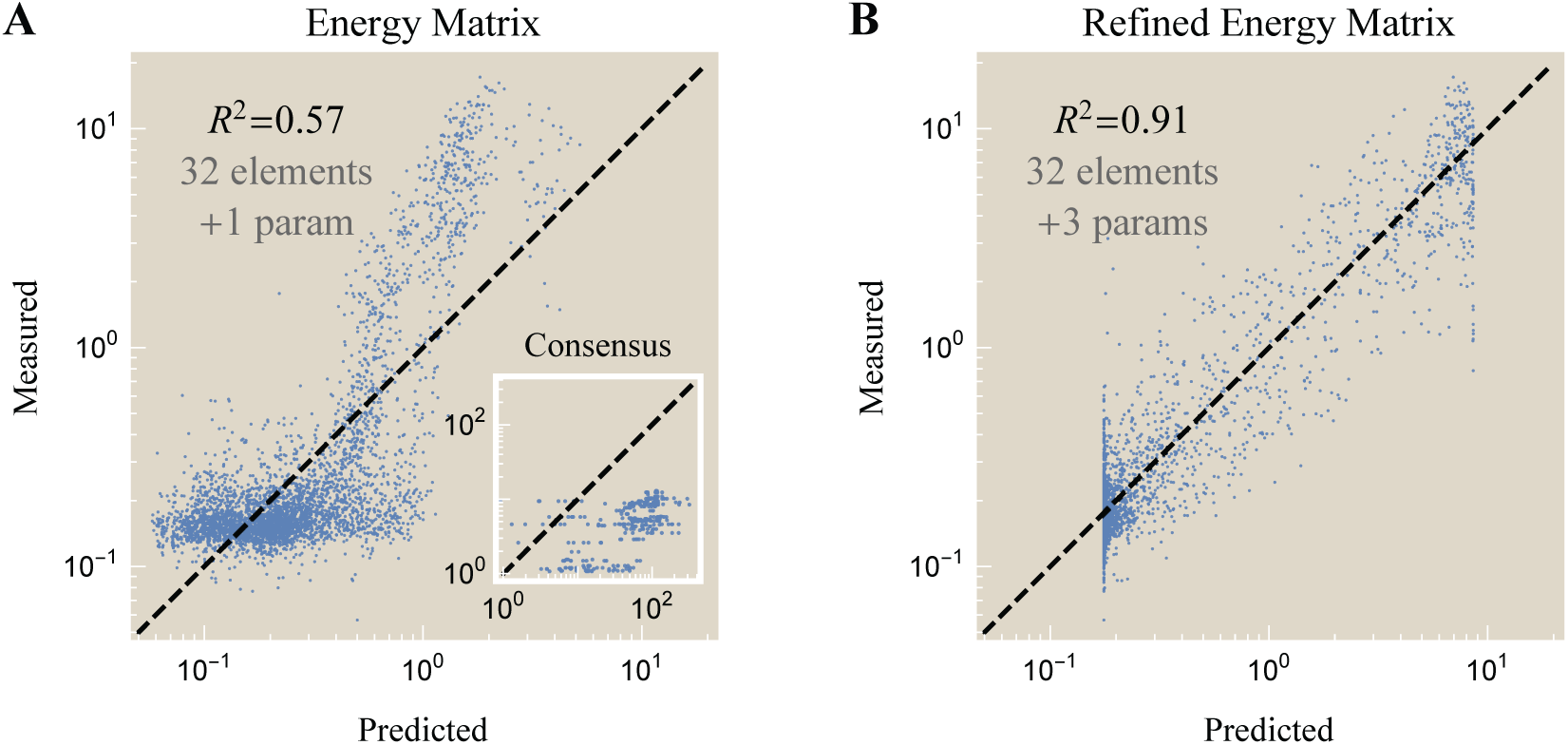
Gene expression of promoters with no UP element. Model predictions using (A) an energy matrix (Eq. 1) where the -35 and -10 elements independently contribute to RNAP binding and (B) a refined energy matrix (Eq. 3) where the two sites contribute cooperatively. Inset: The epistasis-free nature of the energy matrix model makes sharp predictions about the gene expression of the consensus -35 and -10 sequences that markedly disagree with the data. Parameter values given in Appendix B.

Since Eq. 2 still assumes that each promoter element contributes additively and independently to the total RNAP binding energy, it also makes sharp predictions that markedly disagrees with the data (see Appendix C). Inspired by these inconsistencies, we postulated that certain promoter elements, most likely the -35 and -10 sites, may not contribute synergistically to RNAP binding.

To that end, we consider a model for gene expression shown in Fig. 1B where RNAP can separately bind to the -35 and -10 sites. RNAP is assumed to elicit a large level of gene expression *r*_max_ when fully bound but the smaller level *r*_0_ when unbound or partially bound. Importantly, the Boltzmann weight of the fully bound state contains the free energy *E*_int_ representing the avidity of RNAP binding to the -35 and -10 sites. Physically, avidity arises because unbound RNAP binding to either the -35 or -10 sites gains energy but loses entropy, while this singly bound RNAP attaching at the other (-10 or -35) site again gains energy but loses much less entropy, as it was tethered in place rather than floating in solution. Hence we expect 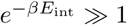, and including this avidity term implies that RNAP no longer binds independently to the -35 and -10 sites.

Our coarse-grained model of gene expression neglects the kinetic details of transcription whereby RNAP transitions from the closed to open complex before initiating transcription. Instead, we assume that there is a separation of timescales between the fast process of RNAP binding/unbinding to the promoter and the other processes that constitute transcription. In the quasi-equilibrium framework shown in Fig. 1B, gene expression is given by the average occupancy of RNAP in each of its states, namely,

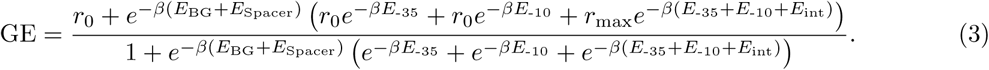

We call this expression a refined energy matrix model since it reduces to the energy matrix Eq. 1 (with constant of proportionality 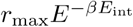) in the limit where gene expression is negligible when the RNAP is not bound (*r*_0_ ≈ 0) and the promoter is sufficiently weak or the RNAP concentration is sufficiently small that polymerase is most often in the unbound state (so that the denominator ≈ 1). The background and spacer are assumed to contribute to RNAP binding in both the partially and fully bound states, an assumption that we rigorously justify in Appendix D.

Fig. 2B demonstrates that the refined energy matrix Eq. 3 is better able to capture the system’s behavior (*R*^2^ = 0.91) while only requiring two more parameters (*r*_0_ and *E*_int_) than the energy matrix model Eq. 1. The sharp boundaries on the left and right represent the minimum and maximum levels of gene expression, *r*_0_ = 0.18 and *r*_max_ = 8.6, respectively (see Appendix E). The refined energy matrix predicts that the top 5% of promoters will exhibit expression levels of 7.6 (compared to 8.5 measured experimentally) while the weakest 5% of promoters should express at 0.2 (compared to the experimentally measured 0.1). In addition, this model quickly gains predictive power, as its coefficient of determination only slightly diminishes (*R*^2^ = 0.86) if the model is trained on only 10% of the data and used to predict the remaining 90%.

### Epistasis-Free Models of Gene Expression Lead to Sharp Predictions that Disagree with the Data

To further validate that the lower coefficient of determination of the energy matrix approach (Eq. 1) was not an artifact of the fitting, we can utilize the epistasis-free nature of this model to predict the gene expression of double mutants from that of single mutants. More precisely, denote the gene expression GE^(0,0)^ of a promoter with the consensus -35 and -10 sequences (and any background or spacer sequence). Let GE^(1,0)^, GE^(0,1)^, and GE^(1,1)^ represent promoters (with this same background and spacer) whose -35/-10 sequences are mutated/consensus, consensus/mutated, and mutated/mutated, respectively, where “mutated” stands for any non-consensus sequence. As derived in Appendix D, the gene expression of these three later sequences can predict the gene expression of the promoter with the consensus -35 and -10 without recourse to fitting, namely,

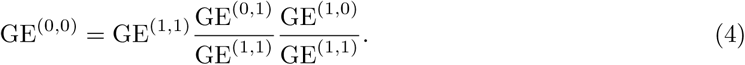

The inset in Fig. 2A compares the epistasis-free predictions (*x*-axis, right-hand side of Eq. 4) with the measured gene expression (*y*-axis, left-hand side of Eq. 4). These results demonstrate that the simple energy matrix formulation fails to capture the interaction between the -35 and -10 binding sites. While this calculation cannot readily generalize to the refined energy matrix model since it exhibits epistasis, it is analytically tractable for weak promoters where the refined energy matrix model displays a marked improvement over the traditional energy matrix model (see Appendix C).

### RNAP Binding to the UP Element occurs Independently of the Other Promoter Elements

Having seen that the refined energy matrix model (Eq. 3) can outperform the traditional energy matrix analysis on promoters with no UP element, we next extend the former model to promoters containing an UP element. Given the importance of the RNAP interactions with the -35 and -10 sites seen above, Fig. 3A shows three possible mechanisms for how the UP element could mediate RNAP binding. For example, the C-terminal could bind strongly and independently so that RNAP has three distinct binding sites. Another possibility is that the RNAP *α*CTD binds if and only if the -35 binding site is bound. A third alternative is that the UP element contributes additively and independently to RNAP binding (analogous to the spacer and background).

**Figure 3.**
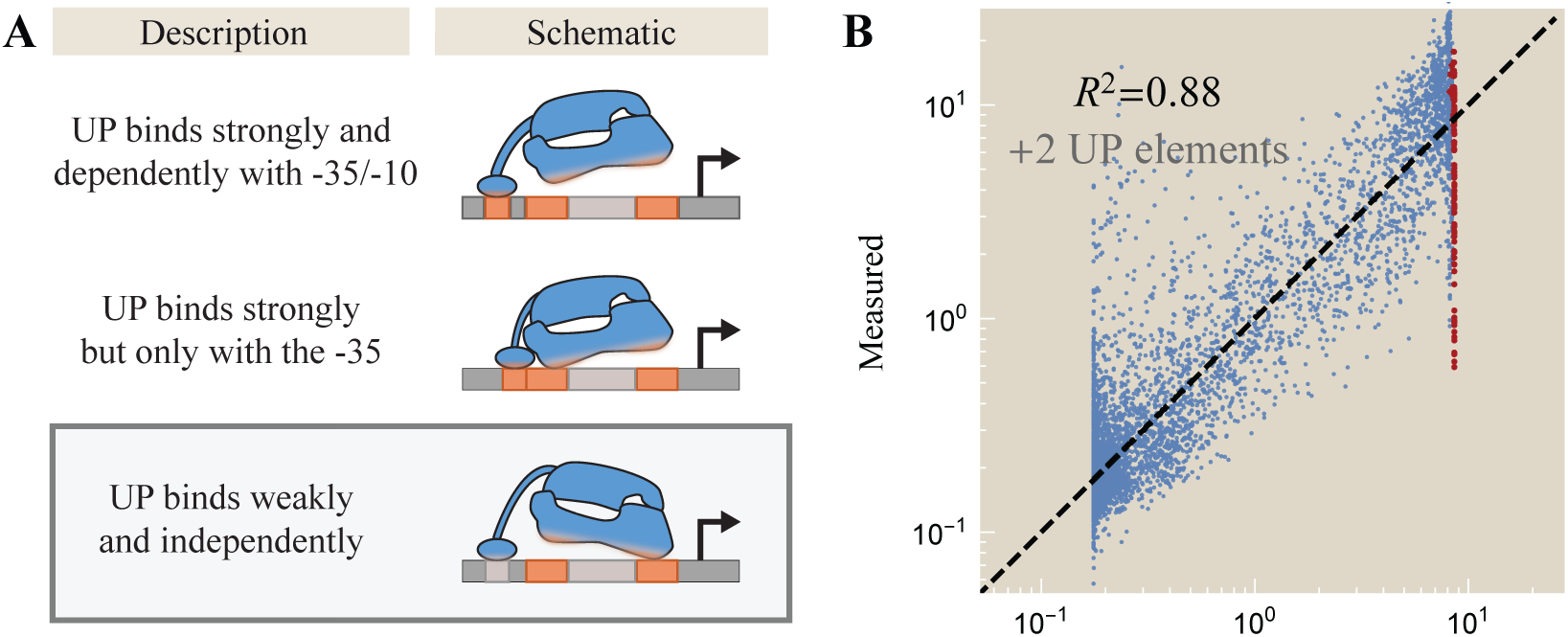
The interaction between RNAP and the UP element. (A) Possible mechanisms by which the RNAP C-terminal can bind to the UP element (orange segments represent strong binding comparable to the -35 and -10 motifs; gray segments represent weak binding comparable to the spacer and background). The data supports the bottom schematic (see Appendix D). (B) The corresponding characterization of 8,192 promoters identical to those shown in Fig. 2 but with one of two UP binding motifs. Red points represent promoters with a consensus -35 and -10. Data was fit using the same parameters as in Fig. 2B and fitting the binding energies of the two UP elements (parameter values in Appendix B).

To distinguish between these possibilities, we analyze the correlations in gene expression between every pair of promoter elements (UP and -35, spacer and background, etc.) to determine the strength of their interaction. Each model in Fig. 3A will have a different signature: The top schematic predicts strong interactions between the -35 and -10, between the UP and -35, and between the UP and -10; the middle schematic would give rise to strong dependence between the -35 and -10 as well as between the UP and -10, while the UP and -35 elements would be perfectly correlated; the bottom schematic suggests that the UP elements will contribute independently of the other promoter elements.

This analysis, which we relegate to Appendix D, demonstrates that the UP element is approximately independent of all other promoter elements (*R*^2^ ≿ 0.6) as are the background and spacer, indicating that the bottom schematic in Fig. 3A characterizes the binding of the UP element. This leads us to the general form of transcriptional regulation by RNAP, shown in Eq. 5.

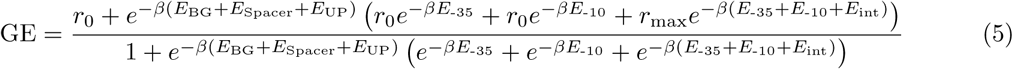

Fig. 3B demonstrates how the expression of all promoters containing one of the two UP elements combined with each of the eight background, spacer, -35, and -10 sequences (2 × 8^4^ = 8,192 promoters) closely matches the model predictions (*R*^2^ = 0.88). Remarkably, since we used the same free energies and gene expression rates from Fig. 2B, characterizing these 8,192 promoters only required two additional parameters (the free energies of the two UP elements). This result emphasizes how understanding each modular component of gene expression can enable us to harness the combinatorial complexity of sequence space.

### Sufficiently Strong RNAP-Promoter Binding Energy can Decrease Gene Expression

Although the 12,288 promoters considered above are well characterized by Eq. 5 on average, the data demonstrate that the full mechanistic picture is more nuanced. For example, Urtecho *et al.* found that gene expression (averaged over all backgrounds and spacers) generally increases for -35/-10 elements closer to the consensus sequences (8). In terms of the gene expression models studied above (Eqs. 1-3), promoters with fewer -35/-10 mutations have more negative free energies *E*_-35_ and *E*_-10_ leading to larger expression. Yet the strongest promoters with the consensus -35/-10 violated this trend, exhibiting *less* expression than promoters one mutation away. Thus, Urtecho *et al.* postulated that past a certain point, promoters that bind RNAP too tightly may inhibit transcription initiation and lead to decreased gene

The promoters with a consensus -35/-10 are shown as red points in Fig. 3B, and indeed these promoters are all predicted to bind tightly to RNAP and hence express at the maximum level *r*_max_ = 8.6, placing them on the right-edge of the data. Yet depending on their UP, background, and spacer, many of these promoters exhibit significantly less gene expression then expected. Motivated by this trend, we posit that the state of transcription initiation can be characterized by a free energy Δ*E*_trans_ relative to unbound RNAP that competes with the free energy Δ*E*_RNAP_ between fully bound and unbound RNAP (see Appendix E).

Assuming the rate of transcription initiation is proportional to the relative Boltzmann weights of these two states, the level of gene expression *r*_max_ in Eq. 5 will be modified to

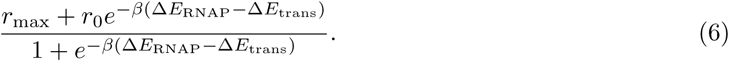

As expected, this expression reduces to *r*_max_ for promoters that weakly bind 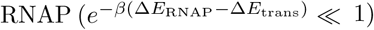 but decreases for strong promoters until it reaches the background level *r*_0_ when the promoter binds so tightly that RNAP is glued in place and unable to initiate transcription. Upon reanalyzing the gene expression data with the inferred value Δ*E*_trans_ = *-*6.2 *k*_*B*_*T* (see Appendix E), we can plot the measured level of gene expression against the predicted RNAP-promoter free energy Δ*E*_RNAP_ as shown in Fig. 4 (stronger promoters to the right). We find that this revised model captures the downwards trend in gene expression observed for the strongest promoters, most of which contain a consensus -35/-10.

**Figure 4.**
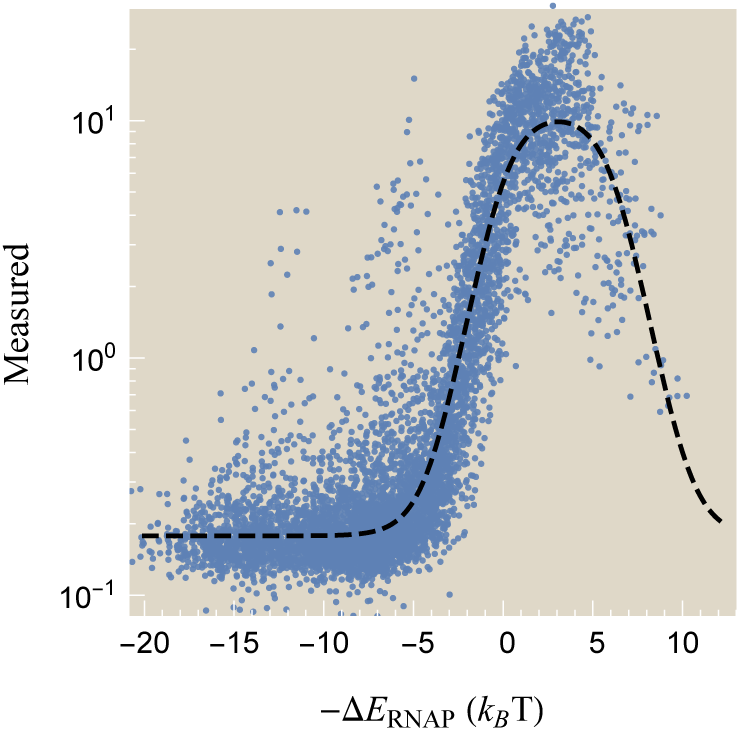
Gene expression is reduced when RNAP binds a promoter too tightly. Measured gene expression versus the inferred promoter strength Δ*E*_RNAP_ relative to the transcription initiation state Δ*E*_trans_ = - 6.2 *k*_*B*_*T* (stronger promoters on the right). The dashed line shows the prediction of the refined energy matrix model.

### The Bivalent Binding of RNAP Buffers its Binding Behavior against Promoter Mutations

In this final section, we investigate how the avidity between the -35 and -10 sites changes the dynamics of RNAP binding. More specifically, we consider the effective dissociation constant governing RNAP binding when both the -35 and -10 sites are intact and compare it to the case where only one site is capable of binding. To simplify this discussion, we focus exclusively on the case of RNAP binding to the -35 and -10 motifs as shown in the rates diagram Fig. 1C, absorbing the effects of the background, spacer, and UP elements into these rates.

At equilibrium, there is no flux between the four RNAP states. We define the effective dissociation constant

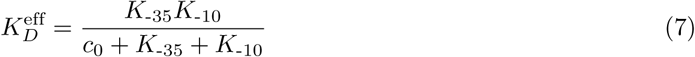

which represents the concentration of RNAP at which there is a 50% likelihood that the promoter is bound (see Appendix F). 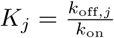 stands for the dissociation constant of free RNAP binding to the site *j* and 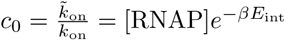 represents the increased local concentration of singly bound RNAP transitioning to the fully bound state (i.e., *E*_int_ and *c*_0_ are the embodiments of avidity in the language of statistical mechanics and thermodynamics, respectively). Note that 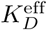 is a sigmoidal function of *K*_-10_with height *K*_-35_ and midpoint at *K*_-10_ = *c*_0_ + *K*_-35_.

Fig. 5 demonstrates how the effective RNAP dissociation constant 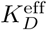 changes when mutations to the -10 binding motif alter its dissociation constant *K*_-10_. When the -35 sequence is weak (dashed lines, *k*_off,-35_ → ∞), 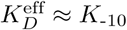 signifying that RNAP binding relies solely on the strength of the -10 site. In the opposite limit where RNAP tightly binds to the -35 sequence (solid lines), the cooperativity *c*_0_ and the dissociation constant *K*_-35_ shift the curve horizontally and bound the effective dissociation constant to 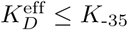. This upper bound may buffer promoters against mutations, since achieving a larger effective dissociation constant would require not only wiping out the -35 site but in addition mutating the -10 site. Finally, in the case where the cooperativity *c*_0_ is large, 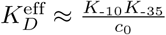 indicating that as soon as one site of the RNAP binds, the other is very likely to also bind, thereby giving rise to the multiplicative dependence on the two *K*_*D*_s.

**Figure 5.**
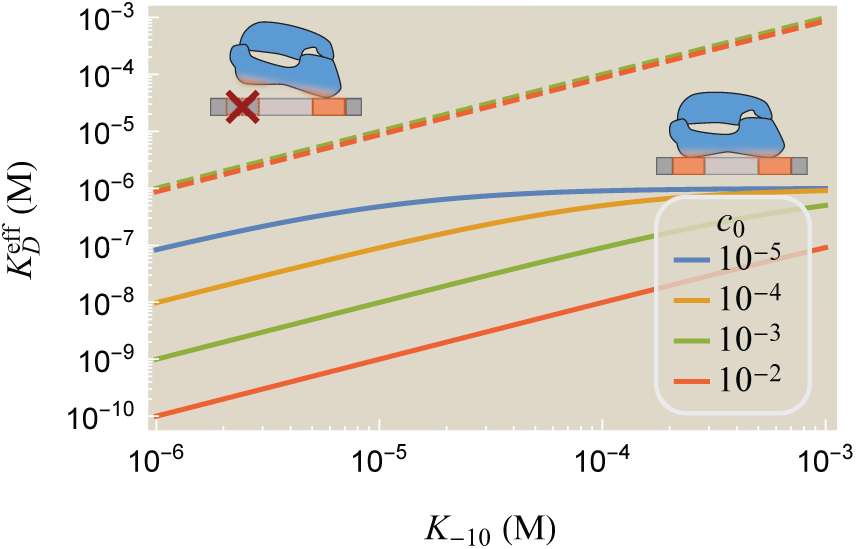
The dissociation between RNAP and the promoter. RNAP binding to a promoter with a strong (solid lines, *K*_-35_ = 1*µ*M) or weak (dashed, *K*_-35_ → ∞) -35 sequence. *c*_0_ represents the local concentration of singly bound RNAP.

To get a sense for how these numbers translate into physiological RNAP dwell times on the promoter, we note that the lifetime of bound RNAP is given by 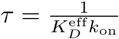 (see Appendix F). Using 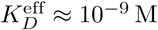 for the strong T7 promoter (15) and assuming a diffusion-limited on-rate 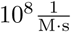 leads to a dwell time of 10 s, comparable to the measured dwell time of RNAP-promoter in the closed complex (16). It would be fascinating if recently developed methods that visualize real-time single-RNAP binding events probed the dwell time of the promoter constructed by Urtecho *et al.* to see how well the predictions of the refined energy matrix model match experimental measurements (16).

## Discussion

While high-throughput methods have enabled us to measure the gene expression of tens of thousands of promoters, they nevertheless only scratch the surface of the full sequence space. A typical promoter composed of 200 bp has 4^200^ variants (more than the number of atoms in the universe). Nevertheless, by understanding the principles governing transcriptional regulation, we can begin to cut away at this daunting complexity to design better promoters.

In this work, we analyzed a recent experiment by Urtecho *et al.* measuring gene expression of over 10,000 promoters in *E. coli* using the *σ*^70^ RNAP holoenzyme (8). These sequences comprised all combinations of a small set of promoter elements, namely, eight -10s, eight -35s, eight spacers, eight backgrounds, and three UPs depicted in Fig. 1A, providing an opportunity to deepen our understanding of how these elements interact and to compare different quantitative models of gene expression.

We first analyzed this data using classic energy matrix models which posit that each promoter element contributes independently to the RNAP-promoter binding energy. As emphasized by Urtecho *et al.* and other groups, such energy matrices poorly characterize gene expression (Fig. 2A, *R*^2^ = 0.57) and offer testable predictions that do not match the data (Appendix C), mandating the need for other approaches (7, 8).

To meet this challenge, we first determined which promoter elements contribute independently to RNAP binding (Appendix D). This process, which was done without recourse to fitting, demonstrated that the -35 and -10 elements bind in a concerted manner that we postulated is caused by avidity. In this context, avidity implies that when RNAP is singly bound to either the -35 or -10 sites, it is much more likely (compared to unbound RNAP) to bind to the other site, similar to the boost in binding seen in bivalent antibodies (17) or multivalent systems (13, 18, 19). Surprisingly, we found that outside the -35/-10 pair, the other components of the promoter contributed independently to RNAP binding.

Using these findings, we developed a refined energy matrix model of gene expression (Eq. 5) that incorporates the avidity of between the -35/-10 sites as well as the independence of the UP/spacer/background interactions. This model was able to characterize the 4,096 promoters with no UP element (Fig. 2B, *R*^2^ = 0.91) and the 8,192 promoters containing an UP element (Fig. 3B, *R*^2^ = 0.88). These results surpass those of the traditional energy matrix model (Fig. 2A, *R*^2^ = 0.57), only requiring two additional parameters that could be experimentally determined (e.g., the interaction energy *E*_int_ arising from the -35/-10 avidity and the level of gene expression *r*_0_ of a promoter with a scrambled -10 motif, a scrambled -35 motif, or with both motifs scrambled).

These promising findings suggest that determining which components bind independently is crucial to properly characterize multivalent systems. It would be fascinating to extend this study to RNAP with other *σ* factors (14) as well as to RNAP mutants with no *α*CTD or that do not bind at the -35 site (20, 21). Our model would predict that polymerases in this last category with at most one strong binding site should conform to a traditional energy matrix approach.

Quantitative frameworks such as the refined energy matrix model explored here can deepen our understanding of the underlying mechanisms governing a system’s behavior. For example, while searching for systematic discrepancies between our model prediction and the gene expression measurements, we found that promoters predicted to have the strongest RNAP affinity did not exhibit the largest levels of gene expression (thus violating a core assumption of nearly all models of gene expression that we know of). This led us to posit a characteristic energy for transcription initiation that reduces the expression of overly strong promoters (Fig. 4). In addition, we explored how having separate binding sites at the -35 and -10 elements buffers RNAP kinetics against mutations; for example, no single mutation can completely eliminate gene expression of a strong promoter with the consensus -35 and -10 sequence, since at least one mutation in both the -35 and -10 motifs would be needed (Fig. 5).

Finally, we end by zooming out from the particular context of transcription regulation and note that multivalent interactions are prevalent in all fields of biology (22), and our work suggests that differentiating between independent and dependent interactions may be key to not only characterizing overall binding affinities but to also understand the dynamics of a system (23). Such formulations may be essential when dissecting the much more complicated interactions in eukaryotic transcription where large complexes bind at multiple DNA loci (24, 25) and more broadly in multivalent scaffolds and materials (12, 13).

## Methods

We trained both the standard and refined energy matrix models on 75% of the data and characterized the predictive power on the remaining 25%, repeating the procedure 10 times. The coefficient of determination *R*^2^ was calculated for *y*_data_ = log_10_(gene expression) to prevent the largest gene expression values from dominating the result. The supplementary Mathematica notebook contains the data analyzed in this work and can recreate all plots.

## Acknowledgements

We thank Suzy Beeler, Vahe Galstyan, Peng (Brian) He, and Zofii Kaczmarek for helpful discussions. This work was supported by the Rosen Center at Caltech and the National Institutes of Health through 1R35 GM118043-01 (MIRA).

## A. The Energy Matrix Model

### A.1 Translating between an Energy Matrix with Base Pair Resolution and Promoter Element Resolution

In this section, we discuss how an energy matrix model with base-pair resolution can be translated into an equivalent model with the resolution of promoter elements. The former model purports that the RNAP-promoter binding energy is composed of independent and linearly additive contributions from each base pair. More precisely, if at position *j* the base *b*_*j*_ (either A, T, C, or G) contributes a free energy 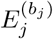 to RNAP binding, then the total free energy of binding is given by 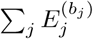 as shown in Fig. S1.

By breaking this sum up over the positions *j* demarking the -35 (*-*35 *≤ j ≤ -*30), spacer (*-*29 *≤ j ≤ -*13), -10 (*-*12 *≤ j ≤ -*7), UP (*-*59 *≤ j ≤ -*38; where “no UP” used a random sequence that did not enhance gene expression), and background (all the remaining base pairs between *-*120 *≤ j <* 30) elements, we achieve an energy matrix model where the free energies *E*_BG_, *E*_-35_, *E*_Spacer_, and *E*_-10_ represent the sum of all base pair contributions of the particular sequence considered. For simplicity, the UP element is not explicitly drawn in the figure.

As shown for two sample sequences in Fig. S1, modifying the -35 sequence while keeping the rest of the promoter unchanged leads to a different *E*_-35_ but keeps *E*_BG_, *E*_Spacer_, and *E*_-10_ unchanged. The expression of the full suite of 12,288 promoters studied in this work can be determined from the free energies of the three UP elements and the eight backgrounds, spacers, -35s, and -10s.

**Figure S1.**
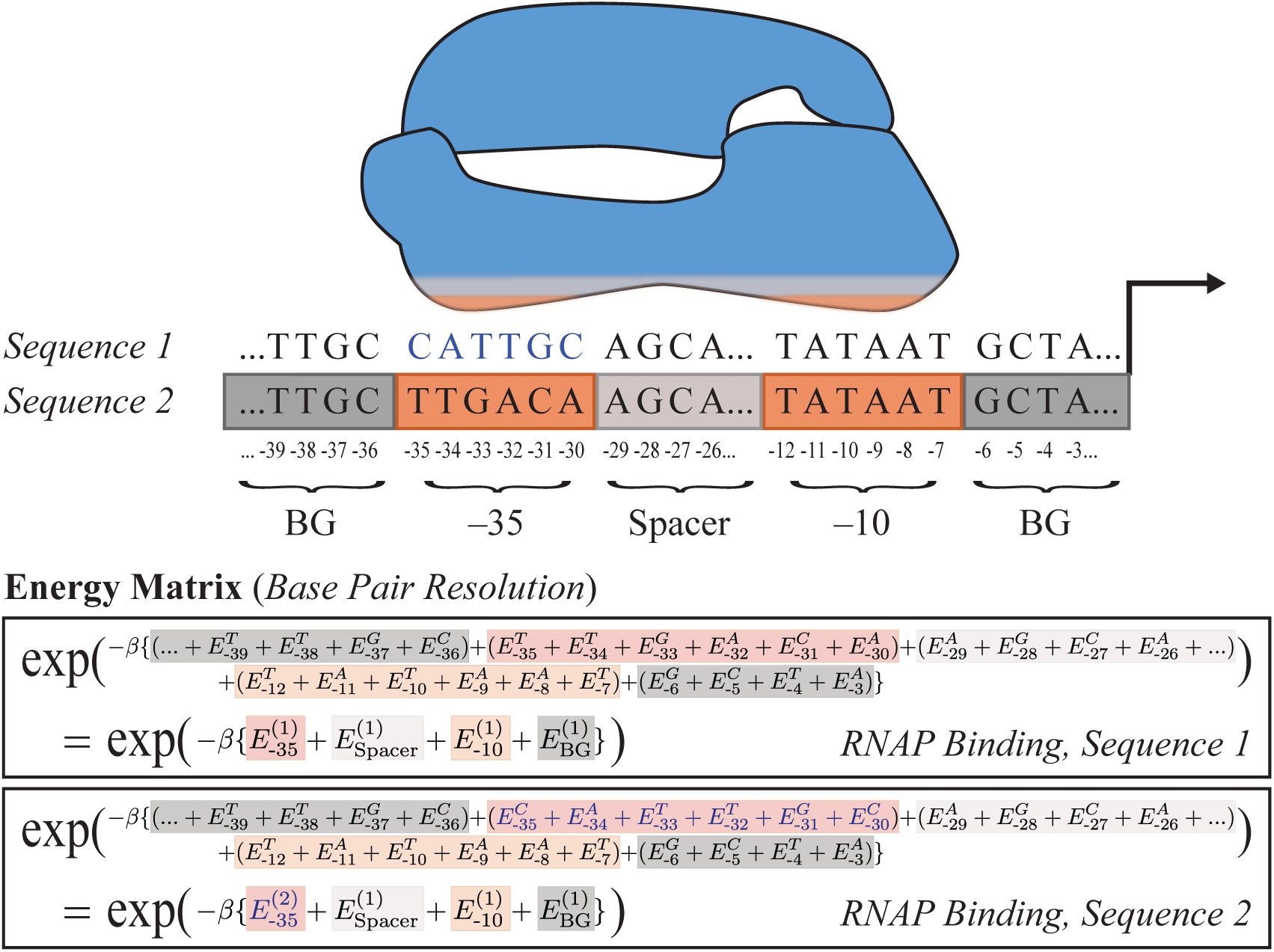
An energy matrix model with base pair resolution translates into an energy matrix model with promoter element resolution. Each promoter element contributes to RNAP binding with free energy given by the sum of its free energies from its base pairs. The two sample sequences shown only differ in their -35 sequence (highlighted blue in Sequence 1), resulting in different values of 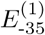 and 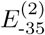 but the same free energies for the remaining promoter elements.

### A.2 Characterizing the Dependence of Gene Expression on RNAP Copy Number

In this section, we explicitly write the dependence of RNAP copy number embedded within the free energies of RNAP binding in Eqs. 1 and 2, thereby making contact with previous models of gene regulation (27). To that end, we consider *P* RNAP molecules that are free to bind anywhere along a bacterial genome with *N*_NS_ <u>n</u>on-specific base pairs (i.e., potential RNAP binding sites outside of our promoter of interest). Let Δ*ϵ* be the average energy difference between RNAP bound to the specific promoter versus at any other location along the genome. By definition, the free energy of RNAP binding considered in this work is given by both the entropic and energetic contributions of this binding, namely,

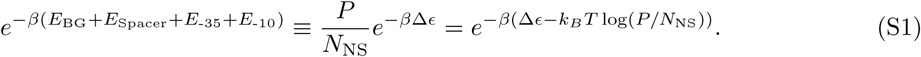

Because the gene expression for each promoter generated by Urtecho *et al.* was measured under the same experimental condition, the RNAP copy number is consistent across all constructs, and hence the constant *k*_*B*_*T* log (*P/N*_NS_) can be absorbed into the free energies. If these measurements were repeated under experimental conditions where the RNAP copy number is halved 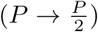, the total free energy of RNAP binding considered in this work would need to be correspondingly modified from (*E*_BG_ + *E*_Spacer_ + *E*_-35_ + *E*_-10_) *→* (*E*_BG_ + *E*_Spacer_ + *E*_-35_ + *E*_-10_ + *k*_*B*_*T* log 2).

## B. Model Fitting and Parameter Values

The energy matrix model (Eq. 1) was solved as a least-squares problem that only fit the promoters in Fig. 2A with no UP element. The refined energy matrix model (Eq. 3) was fit using nonlinear regression on promoter sequences with and without an UP element in order to obtain a single, self-consistent set of parameters capable of capturing the data in Fig. 2B and Fig. 3B. The fitting of both models is presented in the supplementary Mathematica notebook.

The coefficient of determination *R*^2^ was calculated for *y*_data_ = log_10_(gene expression) to prevent the largest gene expression values from dominating the result. We trained both the regular and refined energy matrix models on 75% of the data and characterized the predictive power on the remaining 25%, repeating the procedure 10 times. The exact form used was

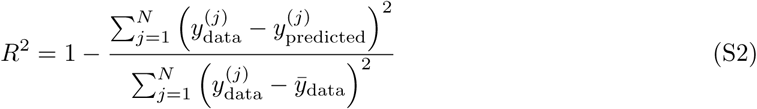

where *y*_predicted_ is the vector of the *N* measurements of log_10_(gene expression) predicted by the model and 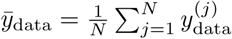 is the average of the logarithmic gene expression data. In this form, the *R*^2^ represents the fraction of variance in the measured gene expression data that arises from the variance in the predicted gene expression data. To test the predictive power of each model, we also trained both models on only 10% of the data and used it to predict the gene expression of the remaining 90% of promoters. We found that the coefficient of determination *R*^2^ only slightly decreased from 0.57 →0.54 for the energy matrix model and from 0.91 → 0.86 for the refined energy matrix model when fitting on this much smaller training set, demonstrating that these models require no more than a thousand promoters to reach their full predictive power.

Table S1 shows the parameter values inferred by the energy matrix (Fig. 2A) and refined energy matrix (Fig. 2B and Fig. 3B) models. Due to the large number of parameters involved, both models exhibit parameter degeneracy (16) where disparate sets of parameters yield nearly identical results. For example, all of the free energies of the spacer elements can be increased by an arbitrary amount provided that the free energies of all background elements are decreased by this same amount (with similar degeneracy holding between other pairs of promoter elements). To circumvent this degeneracy, one -35, one spacer, one -10, one UP, and one background element (denoted by asterisks in Table S1) were fixed to their corresponding value in the energy matrix model, and as such, the parameters below may not represent the binding energies of the promoter elements, but rather only one possible embodiment of these values according to each model.

We point out that our model coarse-grains kinetic details of transcription (e.g., transcription initiation, elongation, transcriptional bursting) into the levels of gene expression *r*_*j*_ shown in Fig. 1B. Modifying the promoter sequence (i.e., considering different spacers or backgrounds) may well change these rates, although our model assumes that such changes only affect the RNAP-promoter binding affinity. If experiments measure the changes in these kinetic rates, they could either be incorporated into the *r*_*j*_ or into an expanded model that explicitly takes these steps of transcription into account (28).

We note from Urtecho *et al.* that the UP elements were named because they increased transcription by 136-fold and 326-fold *in vivo* relative to the physiological *rrnb* P1 UP element. Thus, it follows that the free energy of the 326-fold UP should be smaller than that of the 136-fold UP which should be smaller than the free energy of having no UP element, as seen in Table S1. Additionally, we point out that all spacer elements are 17 bp long; RNAP binding is highly dependent on this length, and promoters with longer or shorter spacers may influence the -35 and -10 binding free energies. Lastly, the sequence composition of all spacers and backgrounds is given in Ref. (8).

**Table S1.**
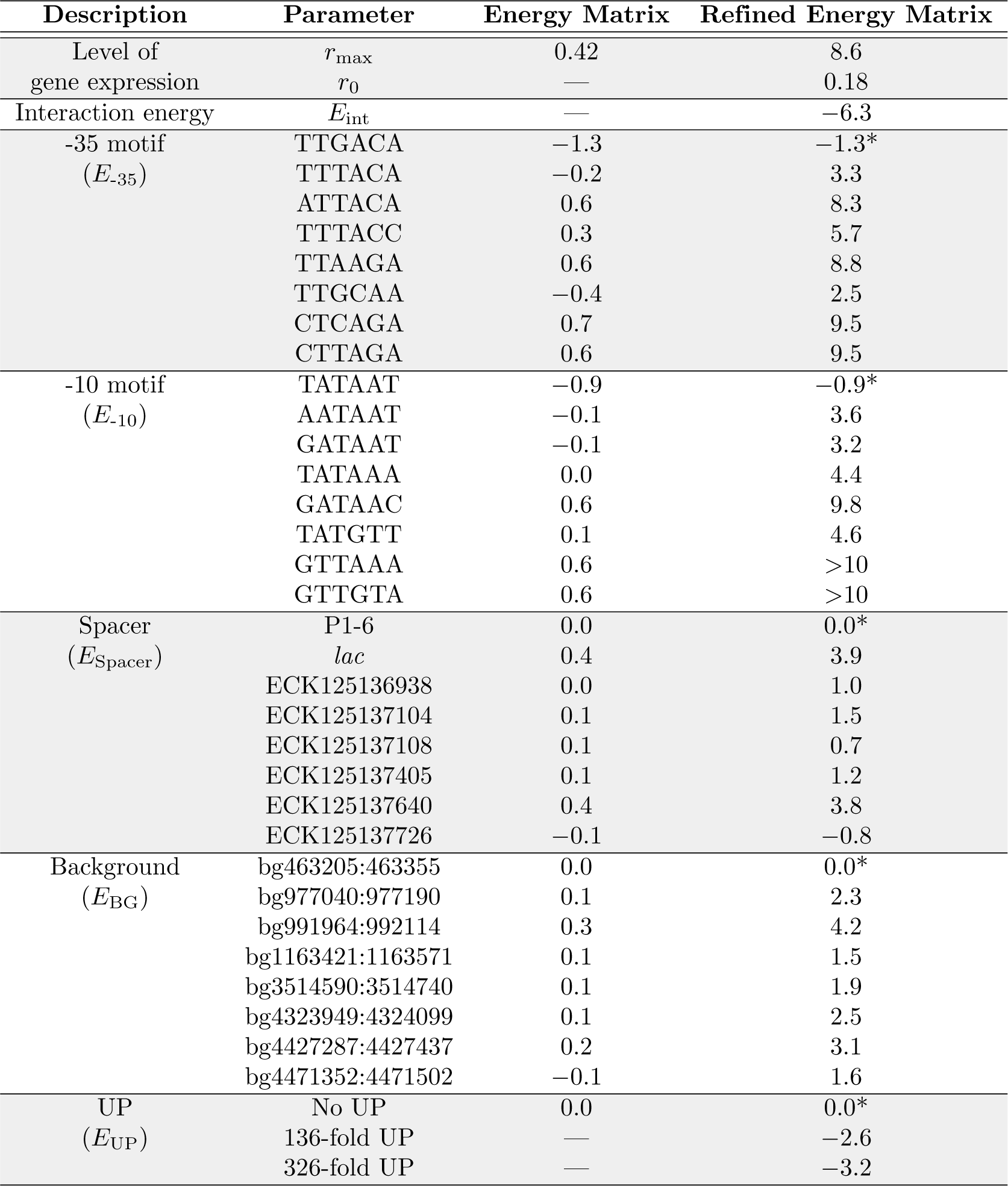
Parameter values for the models of transcriptional regulation considered in this work. The levels of gene expression (*r*_0_ and *r*_max_) are in the same arbitrary units as the experimental measurements (8) while the energies are all in *k*_*B*_*T* units (energies that are more negative indicate tighter binding). The original nomenclature from Table S1 in Ref. (8) is used for each promoter element. Parameter denoted by an asterisk (*) represent values that were fixed to their corresponding value in the energy matrix model to prevent parameter degeneracy.

## C. Comparing the Energy Matrix and Refined Energy Matrix Models of Gene Expression

### C.1 An Epistasis-Free Energy Matrix Model with Saturation does not Capture the Trends in Gene Expression Exhibited by the Data

As shown in Fig. 2A, the simplest model where gene expression is proportional to 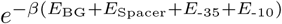 (in the absence of an UP element) fails to characterize the data (*R*^2^ = 0.57). In contrast, the refined energy matrix model in Fig. 2B quantitatively matches the behavior of the spectrum of promoters (*R*^2^ = 0.91). Thus, it behooves us to examine what properties of the latter model are necessary to achieve this concordance with the data.

To that end, we consider an intermediate model where gene expression is given by

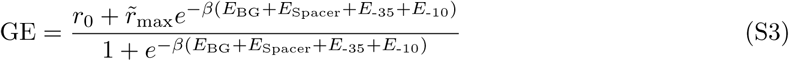

where *r*_0_ represents the minimum level of gene expression in the absence of RNAP binding, 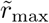 denotes the amount of gene expression when RNAP is fully bound to the promoter, and the *E*_*j*_ represent the free energy contribution of the promoter element *j*. Note that this model represents the limit of a very strong interaction energy in Eq. 3 (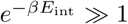 with 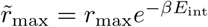) where RNAP is either unbound or fully bound to the promoter.

Fig. S2A demonstrates that the data is well characterized by Eq. S3 (*R*^2^ = 0.91). Therefore, one key feature missing from the simplest energy matrix model description Eq. 1 was that gene expression will saturate once RNAP binding becomes sufficiently strong (or, mathematically, that the denominator 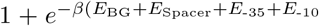 must include the RNAP binding term). Note that the results of this energy matrix model with saturation are nearly identical to the results of the refined energy matrix model in Fig. 2B. Indeed, since the inferred interaction energy *E*_int_ = - 6.3 *k*_*B*_*T* between the -35 and -10 sites is large and negative (see Table S1), it is not surprising that the two models produce similar results for the majority of promoters.

Intuitively, the difference between these two models will emerge in their predictions for promoters with weak expression. As we will show below, the energy matrix model with saturation (Eq. S3) is epistasis-free: given the gene expression of any initial promoter and two mutants of that promoter, we can predict the expression of the double mutant. If, for example, the initial promoter exhibits weak gene expression and the two mutants exhibit a medium level of gene expression, then the double mutant would be predicted to exhibit a large amount gene expression. As will be explained below, the resulting predictions shown in Fig. S2B are highly damning. On the other hand the refined energy matrix model (Eq. 3) predicts a more complex relationship between these four promoters, and in the Appendix C.2 we examine an analytically tractable limit to show that this model better recapitulates the gene expression measurements.

We proceed by utilizing the epistasis-free nature of Eq. S3. A key feature of the following analysis is that it will not require any model fitting, and hence for the remainder of this Appendix we proceed as if we have no knowledge of the parameter values in Table S1. To begin, we approximate the values of *r*_0_ ≈ 0.2 and 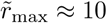 from the gene expression data (the minimum and maximum *y*-values in Fig. S2A, averaging by eye to account for noise). These two values, together with the gene expression measurements for every construct, will be sufficient to make our epistasis-free predictions without explicitly determining any of the *E*_*j*_.

As in the main text, denote the gene expression GE^(0,0)^ of a promoter with the consensus -35 and-10 sequences (and any background or spacer sequence). Let GE^(1,0)^, GE^(0,1)^, and GE^(1,1)^ represent promoters (with this same background and spacer) whose -35/-10 sequences are mutated/consensus, consensus/mutated, and mutated/mutated, respectively. Eq. S3 can be inverted to determine

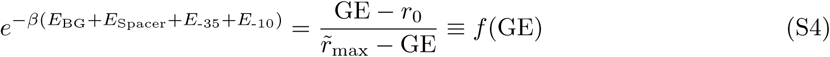

for the double mutant with GE^(1,1)^ as well as the two singly mutated promoters with GE^(1,0)^ and GE^(0,1)^, where we have defined the function *f* for convenience. Importantly, since the -35 and -10 binding energies additively and independently contribute to the RNAP-promoter free energy, the left-hand side of Eq. S4 for the unmutated construct is given by 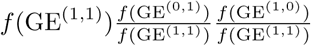 (exactly analogous to Eq. 4 for the simple energy matrix model). Therefore, its gene expression is predicted to be

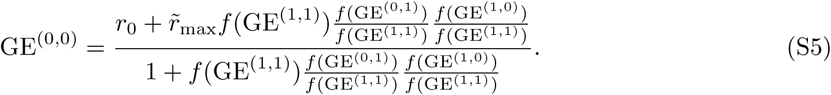

**Figure S2.**
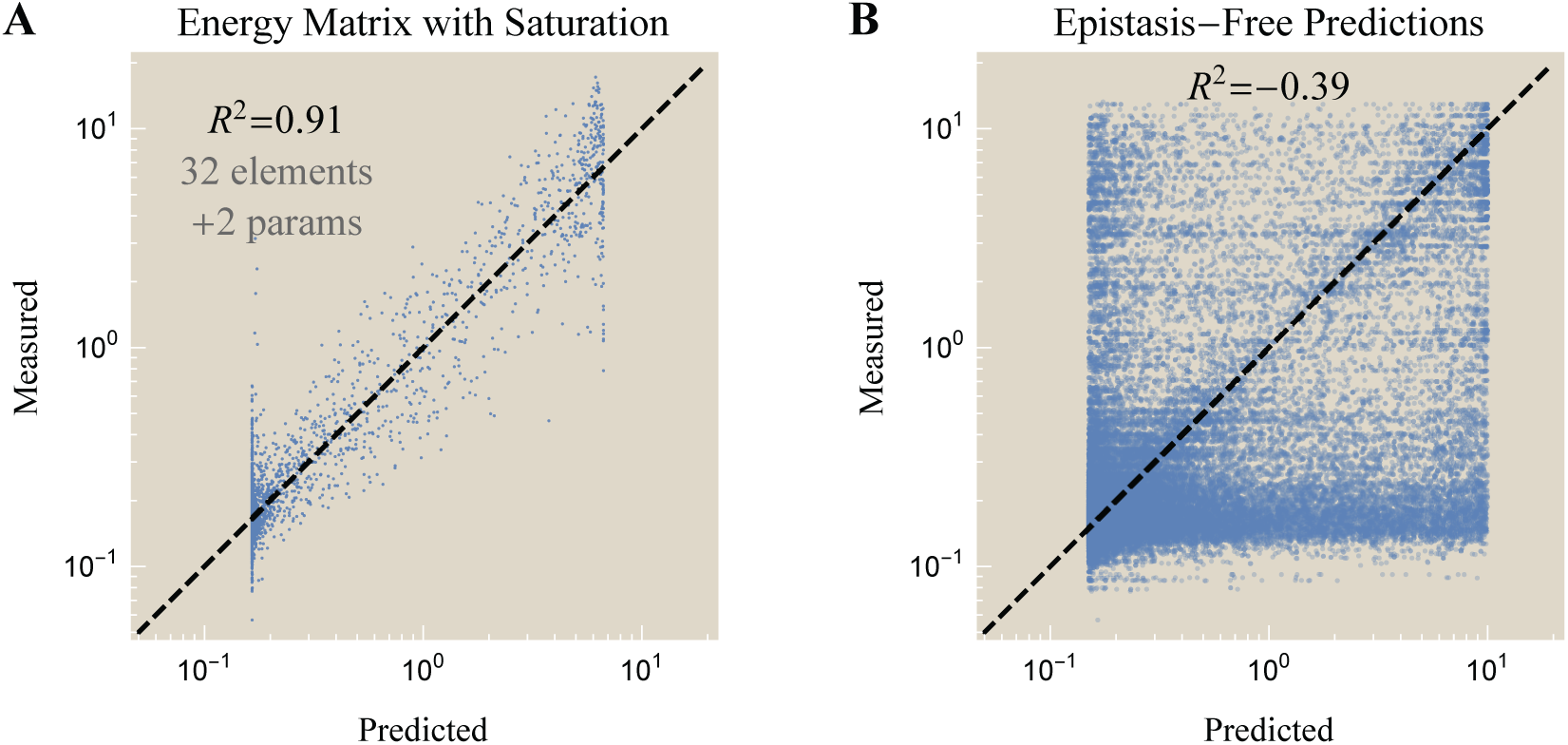
Gene expression represented by an energy matrix model with saturation. (A) Characterization of the same promoters as in Fig. 2 using the energy matrix model with saturation (Eq. S3) with essentially identical fit quality as the refined energy matrix. (B) Since this model assumes that the RNAP-promoter binding energy is epistasis-free (with the -35 and -10 binding sites contributing additively and independently to the RNAP binding energy), the gene expression of double mutants can be predicted from the expression of single mutants without resorting to fitting (Eqs. S4 and S5). The large deviations demonstrate that the energy matrix with saturation cannot characterize the gene expression of these constructs.

Fig. S2B shows the results of these epistasis-free predictions. Because Eq. S5 applies to *any* pairs of -35 and -10 elements with the same BG and spacer, there is a combinatorial explosion of predictions, providing a solid test for this model. As can be seen, aside from the plethora of data points with correctly-predicted low gene expression in the bottom-left corner of the plot, there are large swathes of data points that do not fall on the expected diagonal line, indicating that the epistasis-free prediction in Eq. S3 cannot accurately capture the gene expression of the constructs considered here. In the next section, we show that the refined energy matrix is better equipped to characterize these cases. Notably, these results indicate that although a model may fit the majority of data on average (as in Fig. S2A), it may nevertheless make spurious predictions. Such hidden gems may go unnoticed when a pure-fitting mentality is applied to the wealth of data that is becoming increasingly easy to generate.

### C.2 A Refined Energy Matrix outperforms the Energy Matrix Model in the Limit of a Weak -35 or Weak -10 RNAP Binding Site

In section C.1, we showed that an energy matrix model with saturation Eq. S3 is epistasis-free and hence makes sharp predictions that are inconsistent with the data (Fig. S2B). In this section, we consider the refined energy matrix model Eq. 3 where binding to the -35 and -10 sites is no longer independent. Because this latter model exhibits epistasis, we will restrict our analysis to the limit of weak promoters with no UP element where we can approximate the refined energy model and compare its results to the energy matrix model with saturation. As before, we proceed without referencing the parameter values in Table S1 to emphasize that this analysis can be done without recourse to fitting.

We define GE^(1,1)^, GE^(1,0)^, GE^(0,1)^, and GE^(0,0)^ as in section C.1, but we will restrict our attention to promoters where the original sequence exhibits low gene expression (GE^(1,1)^ 0.25) and the two mutants exhibit medium gene expression (0.25 ≾ GE^(1,0)^, GE^(0,1)^ ≿ 1.0). For such cases, we expect that the predicted gene expression GE^(0,0)^ of the double mutant will be larger in the refined energy matrix model (Eq. 3) than the energy matrix model with saturation (Eq. S3) due to the avidity between the -35 and -10 sites. In other words, the refined energy matrix model acknowledges that the -35 and -10 sites bolster each other and consequently predicts larger gene expression when both sites exhibit even a moderate capability of binding.

As discussed in section C.1, GE^(0,0)^ is exactly given by Eq. S5 in the energy matrix model with saturation. Applying that result to the present case of weak promoters (GE^(1,1)^ 0.25) with medium-strength singly mutants (0.25 ≾ GE^(1,0)^, GE^(0,1)^ ≿ 1.0), Fig. S3A shows that this model generally underestimates the gene expression of these promoters. This serves as a promising indicator that the avidity of RNAP binding is missing from such an approach.

We next turn to the more complex refined energy matrix framework. Because this model exhibits epistasis, the relationship between gene expression is more complex and hence we only roughly approximate GE^(0,0)^. To that end, it behooves us to generalize the levels of gene expression in Fig. 1(B) so that RNAP bound only at the -35 site leads to a gene expression level of *r*_-35_ while RNAP bound only at the -10 site elicits *r*_-10_ gene expression (satisfying *r*_0_ *< r*_-35_, *r*_-10_ *≤ r*_max_), leading to

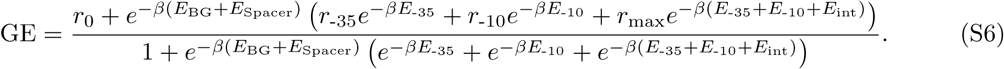

In the main text, we assumed that *r*_-35_ = *r*_-10_ = *r*_0_ for simplicity (and because relaxing this assumption does not qualitatively change any of our results). Here, we will keep these more general rates, as it will aid in the following analysis.

Using Eq. S6, we can approximate gene expression for our four promoters of interest,

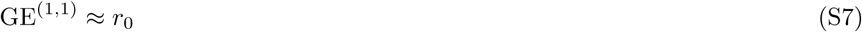

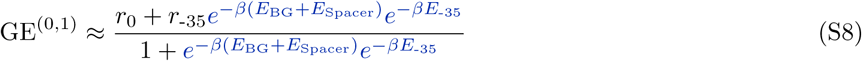

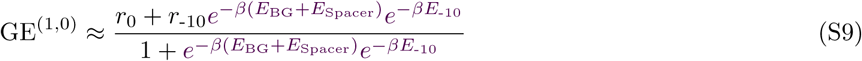

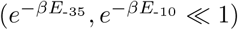

In Eq. S7, we used the fact that the promoter is very weak (GE^(1,1)^ ≲ 0.25) to infer that RNAP is unable to bind at either the -35 or -10 sites 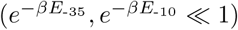 . Since replacing the -35 site slightly improves gene expression (0.25 ≲ GE^(0,1)^ ≲ 1.0), we only keep the -35 binding term in Eq. S8 but continue to neglect the -10 terms (assuming that binding to the -10 is sufficiently unfavored that it overwhelms the avidity term, 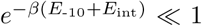. Analogous statements hold for GE^(1,0)^ and the -10 site in Eq. S9. Lastly, when both the -35 and -10 sites are replaced (Eq. S10), the fully bound RNAP state will dominate over the two partially bound states due to avidity.

For every set of four promoters satisfying our criteria, we can use Eq. S7 to infer *r*_0_, Eq. S8 to solve for 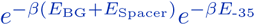 (in terms of *r*_-35_), and Eq. S9 to solve for 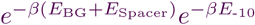 (in terms of *r*_-10_). In addition, we can directly estimate *r*_max_ ≈10 directly from the maximum gene expression of all promoters. Combining these statements, we can rewrite Eq. S10 as

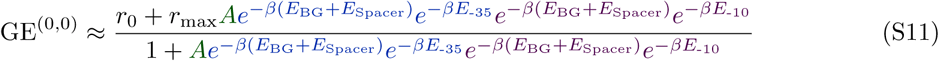

with the unknown quantity 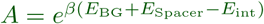. Therefore, the three unknown constants *r*_-35_, *r*_-10_, and *A* would permit us to predict GE^(0,0)^ using single and double mutant data within the refined energy matrix model. To facilitate this, we coarsely approximate that the partially bound RNAP states give rise to intermediate expression levels 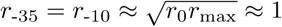 and that the average energy of a background and spacer sequence is negligible compared to the favorable interaction energy which is on the order of several *k*_*B*_*T* leading to 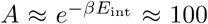. Fig. S3B demonstrates that the refined energy matrix predicts larger gene expression often closer to *r*_max_ ≈10. Although this approximate result for the refined energy matrix model exhibits scatter about the predicted diagonal line, it nevertheless show a marked improvement over the energy matrix model, supporting the notion that avidity is a key concept when predicting the gene expression of mutations that greatly weaken or greatly strengthen the -35 and -10 sites.

**Figure S3.**
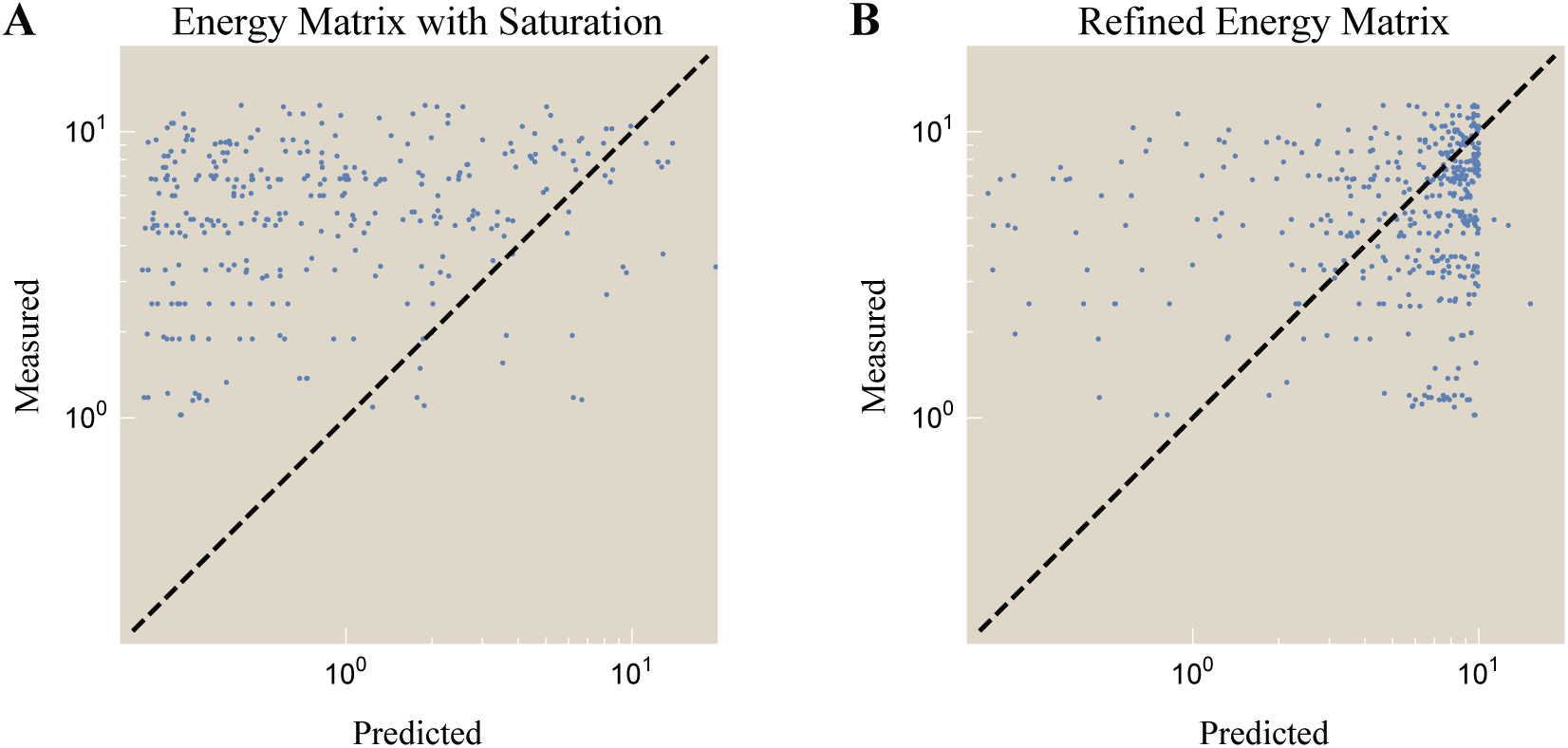
Relating gene expression measurements with minimal fitting. Using gene expression measurements for a weak promoter and two single mutants with higher gene expression, we can predict the expression of the double mutant and compare it to data. (A) The epistasis-free energy matrix model with saturation (Eq. S3) underestimates the gene expression, suggesting that the avidity between the -35 and -10 sites is missing from this analysis. (B) The refined energy matrix model Eq. 3 predicts higher gene expression levels that better characterize the data.

## D. Interactions Between the Different Promoter Elements

In this section, we extend the analysis shown in the Fig. 2A inset to determine the strength of interactions between every pair of promoter elements as shown in Fig. S5A. As an example, Fig. S4 considers the combinations of a promoter with two possible -35 motifs (-35^(1)^ or -35^(2)^) and two possible spacers (Spacer^(1)^ or Spacer^(2)^) with the same UP, -10, and background sequences.

Suppose that the -35 and spacer elements contribute independently to gene expression (GE) so that we can write GE = *f*_1_(*E*_*-*35_)*f*_2_(*E*_Spacer_) as the product of two functions *f*_1_ and *f*_2_ (in the standard energy matrix model, *f*_1_(*E*) = *f*_2_(*E*) = *e*^*-βE*^). This independence implies that the system has no epistasis, namely,

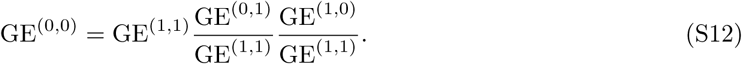

Thus, for all possible pairs of -35 and spacer elements, we can compare the predicted gene expression given by Eq. S12 with the experimental measurements to discern whether these two segments of the promoter contribute independently to gene expression. In the following analysis, we will also restrict ourselves to promoters where GE *>* 10^*-*0.5^ for all four mutants to ensure that the measurements are within the dynamic range of the experiment (so that we can be certain we are analyzing gene expression measurements and not noise).

### D.1 Characterizing Promoters with no UP Element

We first carry out this analysis on the 4,096 promoters with no UP elements as shown in Fig. S5. In each plot, we compare the epistasis-free predicted GE (*x*-axis) with the measured value (*y*-axis). If two promoter elements independently contribute to gene expression, their data should fall onto the straight line *y* = *x*. We can quantify all deviations from such lines using the coefficient of determination *R*^2^, with smaller *R*^2^ values signifying that the promoter elements are not multiplicatively independent.

This analysis shows that while the -35 and -10 elements interact in a fashion discordant with an energy matrix formulation (leading to a negative *R*^2^), the remaining promoter elements interact approximately independently of each other and can be approximated using an energy matrix model. This rigorously justified our sole consideration of the -35 and -10 binding sites in Fig. 1B, allowing us to avoid, for example, enumerating states where the RNAP is solely bound to the spacer or the background. Instead, the promoter is well approximated by treating the -35 and -10 motifs as cooperative binding sites while the spacer and background contribute independently to RNAP binding (as per Eq. 3).

**Figure S4.**
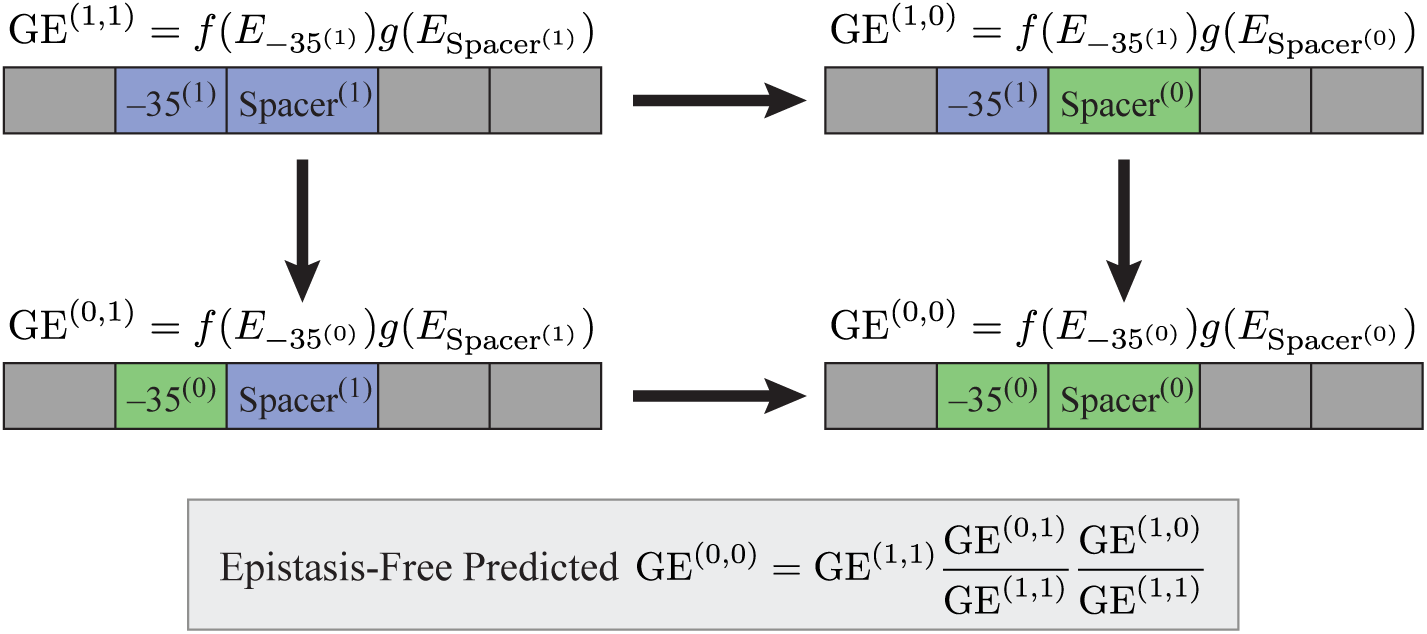
Quantifying the interactions between promoter elements. If the -35 and spacer promoter elements independently contribute to gene expression, then an epistasis-free prediction of gene expression for the double mutant (bottom right) can be predicted using the gene expression of the other three promoters.

**Figure S5.**
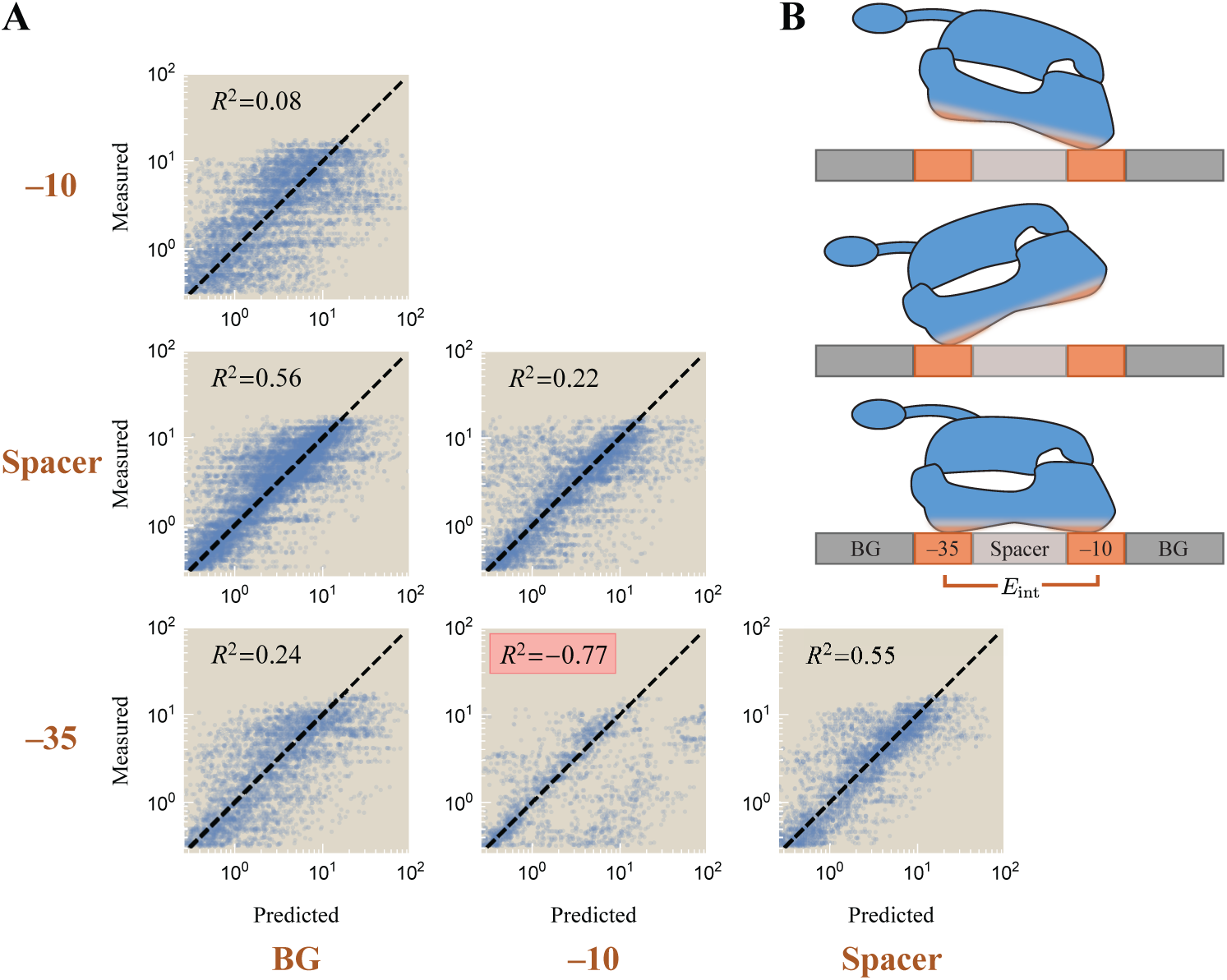
Interactions between the promoter elements with no UP binding site. (A) For every pair of elements (brown labels on the left and bottom), the measured gene expression (*y*-axis) is compared to the epistasis-free prediction (*x*-axis, Eq. S12) assuming that the two promoter elements are independent. Deviations between the predictions and measurements indicate that the two promoter elements interact. Data is plotted with low opacity to better show the general trend across the promoters. (B) The resulting schematic of a promoter with no UP element is that RNAP can bind to either the -35 or -10 sites independently with an avidity interaction when both are bound; the spacer and background (BG) contribute independently to the RNAP binding energy provided RNAP is bound to either the -35 or -10 element.

Lastly, we note that the refined energy matrix model (Eq. 3) does not strictly exhibit the multiplicative independence between the -35 and spacer elements (or any of the other weakly interacting promoter elements) that would lead to an *R*^2^ = 1 expectation, but as we now show it closely approximates multiplicative independence. First, note that using the parameter values from Table S1, the denominator in the refined energy matrix model is approximately 1 because 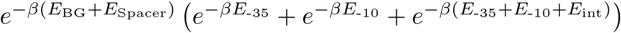 is ≾ 1 for approximately 90% of the promoters. Additionally, the numerator in the model may be dominated by either of its terms: (1) For weak promoters that exhibit low levels of gene expression, 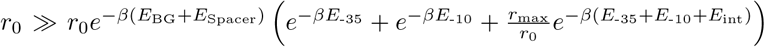 and GE ≈*r*_0_. In the opposite limit where expression is large, 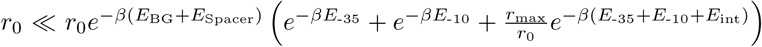 and we can approximate Eq. 5 as

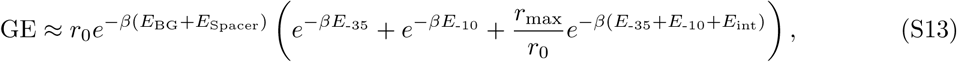

which exhibits the multiplicative independence implied by Eq. S12 between the weakly interacting promoter elements. Practically speaking, this means that in creating Fig. S5, we only considered data points where gene expression was above the background level that we inferred to be 10^*-*0.5^ based on the gene expression measurements in Fig. 2. In summation, Eq. 5 exhibits approximate independence between the weakly interacting promoter elements which can be identified as the plots for which *R*^2^ *>* 0 in Fig. S5.

### D.2 Characterizing Promoters with an UP Element

Here, we extend the analysis in the previous section to a promoter that includes an UP element. As before, we seek to understand whether the UP, -35, spacer, -10, and background elements act independently of each other or whether they interact with avidity to facilitate RNAP binding.

Fig. S6 carries out this analysis using all 12,288 sequences from Urtecho *et al.* for every pair of promoter elements (8). As in the previous section, we find that the -35 and -10 sites do not interact independently (as shown by a negative *R*^2^). We acknowledge that several additional pairs of elements (i.e., -10/BG, -35/BG, and -10/Spacer) exhibit low *R*^2^ values which may arise because: (*i*) Our model only approximately obeys multiplicative independence as discussed in Appendix D.1 (so that *R*^2^≈1 even in the absence of experimental noise) or (*ii*) there may be additional interactions between promoter elements that we neglect, such as the importance of the discriminator (14) or weak RNAP binding sites in the background sequences (5). We proceed by only considering interactions sufficiently strong to induce a negative *R*^2^ value, namely, the avidity between the -35 and -10 motifs, with our eyes wide open to the possibility that more complex models could attempt to capture the full suite of higher-order interactions.

We end this section by analyzing which of the three schematics shown in Fig. 3A best characterizes the binding of the UP element. We note that the UP element appears to be particularly independent (0.6 ≾ *R*^2^) compared all other pairings of elements (0.1 ≾ *R*^2^ ≿0.6), suggesting that the RNAP C-terminal binds weakly provided that either the -35 or -10 motifs are bound (Fig. S6B). This supports the bottom schematic in Fig. 3 and gives rise to the form of gene expression Eq. 5 used in the main text.

To complete this argument, we further note that the middle schematic in Fig. 3A would imply that the UP element only binds when the -35 element is bound, which would result in gene expression of the form

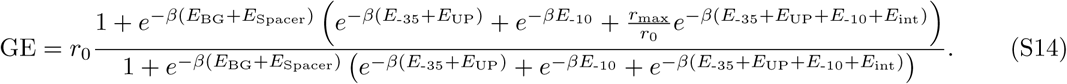

In this case, we would expect a low *R*^2^ between the -35 and -10 elements as well as between the UP and-10 elements, but we would have an *R*^2^ ≈1 between UP and -35 since binding of one forces the binding of the other in this model. Given the larger-than-expected *R*^2^ = 0.56 value between the UP and -10 elements and the smaller-than-expected *R*^2^ = 0.62 value between the UP and -35 elements, this model is unlikely to be correct.

Finally, the top schematic in Fig. 3A implies a low *R*^2^ value between the -35 and -10, the UP and -35, and the UP and -10 elements. In this case, all three elements bind strongly and in a highly dependent manner, so that eight RNAP states would need to be considered (with avidity terms between every pair of elements). Because the *R*^2^ values between the UP/-35 and UP/-10 are larger than expected, this model does not appear to properly characterize RNAP binding, leading us to favor the bottom schematic in Fig. 3A.

**Figure S6.**
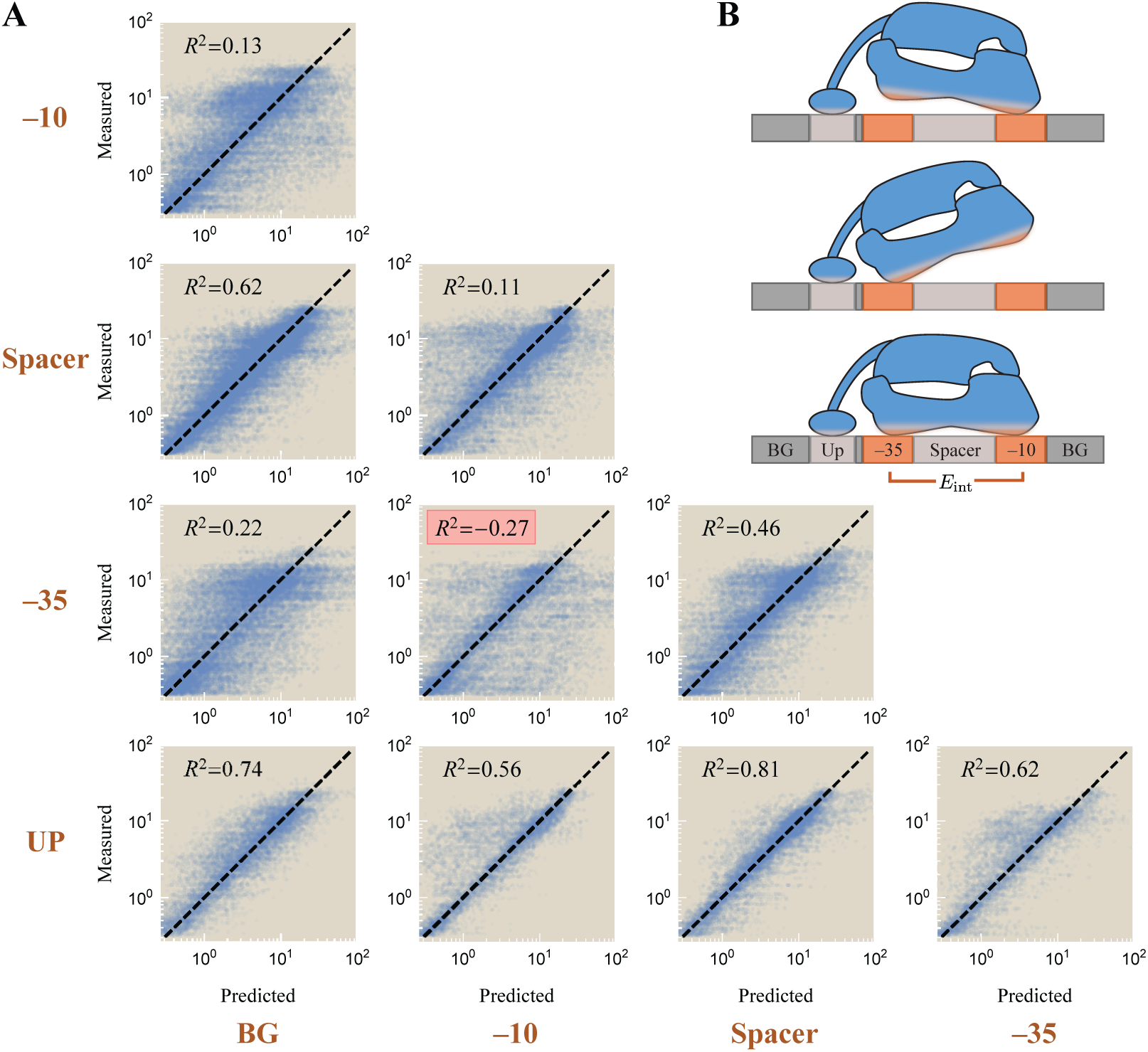
Interactions between the promoter elements with an UP binding site. (A) For every pair of elements (brown labels on the left and bottom), the measured gene expression (*y*-axis) is compared to the epistasis-free prediction (*x*-axis, Eq. S12) assuming that the two promoter elements are independent. (B) The resulting schematic of gene expression where RNAP can bind to either the -35 or -10 sites independently with an avidity interaction when both are bound; the UP, spacer, and background (BG) contribute independently to the RNAP binding energy provided RNAP is bound to either the -35 or -10 element.

## E. RNAP Binding Too Tightly Decreases Gene Expression

All of the gene expression models examined in this work assert that gene expression monotonically increases with the RNAP-promoter binding affinity. In contrast, Urtecho *et al.* found that this monotonic relationship did not hold for the strongest promoters. In other words, gene expression increased as the -35 and -10 motifs approached their consensus sequences (which bind the tightest to RNAP) *except* that promoters with both a consensus -35 and consensus -10 sequence exhibited lower gene expression than the corresponding sequences with one mutation in either motif (8). This suggests that past a certain point, increasing the RNAP-promoter binding energy causes RNA polymerase to bind top tightly, thereby inhibiting gene expression.

In this Appendix, we explore this phenomenon and develop a model to account for it. More specifically, our model will posit that the state of transcription initiation can be characterized by a free energy so that the probability of initiating transcription versus remaining bound on the promoter is given by the Boltzmann weight of the two states.

To start our analysis, Fig. S7A shows the predicted versus measured gene expression of the refined energy matrix model (Eq. 5) for promoters with an UP element. The sharp left edge of the data (gray ellipse) is set by the background level of gene expression *r*_0_ = 0.18 in the absence of RNAP, while the scatter in the *y*-direction can be attributed to noise (since approximately 4,600 promoters have gene expression *<* 0.3, we expect approximately 10 to be 3*σ* away from the predicted value). Meanwhile, the sharp edge on the right side of the data (red ellipse) is caused by promoters with the maximum gene expression *r*_max_ = 8.6, but the downwards scatter is greater than statistically expected (only 450 promoters have gene expression *>* 10, yet there are the same number of outliers 3*σ* away from the mean). This suggests that there is a mechanistic explanation for why these promoters, which the model predicts should bind very tightly to RNAP, exhibit low gene expression.

We next analyzed whether any promoters increased expression when their -35 or -10 sites were replaced by the consensus sequences, but exhibited decreased gene expression when both the -35 and -10 sites became the consensus sequences. Out of the 12,000 constructs, 850 exhibited this pattern of expression. One possible explanation is that although strong binding helps recruit RNAP to the promoter, overly strong binding could inhibit transcription initiation and decrease gene expression. We note, however, that this is a coarse-grained effective model that neglects molecular details of the transition from the closed to open complex, transcription initiation, and other critical steps of RNAP functioning (29). Nevertheless, in the refined energy matrix model, the effect of overly strong RNAP-promoter binding must be to decrease the single parameter *r*_max_ (which represents the level of gene expression when RNAP is fully bound to the promoter), since no other parameters should depend upon the total RNAP-promoter binding strength.

To proceed, we assume that fully bound RNAP with free energy

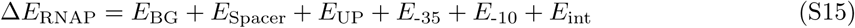

relative to the unbound state can initiate transcription by moving into a transcription initiation state with free energy Δ*E*_trans_ relative to the unbound state as shown schematically in Fig. S7B. Intuitively, bound RNAP will always immediately transcribe when Δ*E*_trans_ Δ-*E*_RNAP_ is large and negative, but when the affinity between the RNAP and promoter becomes sufficiently strong (the case depicted in Fig. S7B), RNAP will prefer to stay bound to the promoter and not transcribe immediately. We posit that the rate of entering the transcribing state, and hence the rate of gene expression *r*_max_ in Fig. 1B, should be modified to

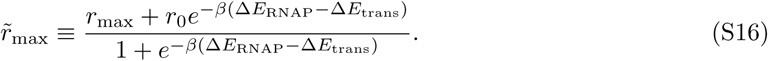

For promoters whose RNAP binding is far weaker than transcription initiation 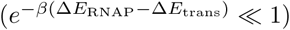, this rate reduces to the constant value *r*_max_. Increasing the RNAP-promoter affinity decreases Δ*E*_RNAP_ which leads to a decrease in the level of gene expression. In the limit of an infinitely strong promoter (Δ*E*_RNAP_ *→ −∞*), RNAP is glued in place and unable to transcribe, thereby reducing the level of gene expression to the background level *r*_0_

Fig. S7C shows the gene expression data refit to the refined energy matrix model with the maximal level of gene expression given by Eq. S16 (using Δ*E*_trans_ = -6.2 *k*_*B*_*T* inferred by nonlinear regression). We note that using this model eliminates the sharp right edge of the data (red ellipse in Panel A), signifying that the promoters with extremely tight RNAP binding have shifted left, moving closer to the level of gene expression predicted by the model. Fig. S7D compares the predicted −Δ*E*_trans_ for each promoter (using the best fit parameter in Table S1) against the measured level of gene expression. To facilitate a comparison with the refined energy matrix model, we overlay this data with the approximate predicted level of gene expression

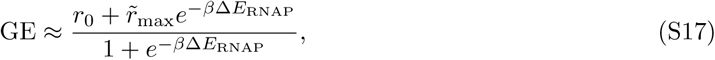

where we have ignored the two partially bound RNAP states and used the maximum level of gene expression in Eq. S16. Although only a small number of promoters exhibits sufficiently strong binding that diminishes their gene expression, the data exhibits a clear downwards trend in this limit.

**Figure S7.**
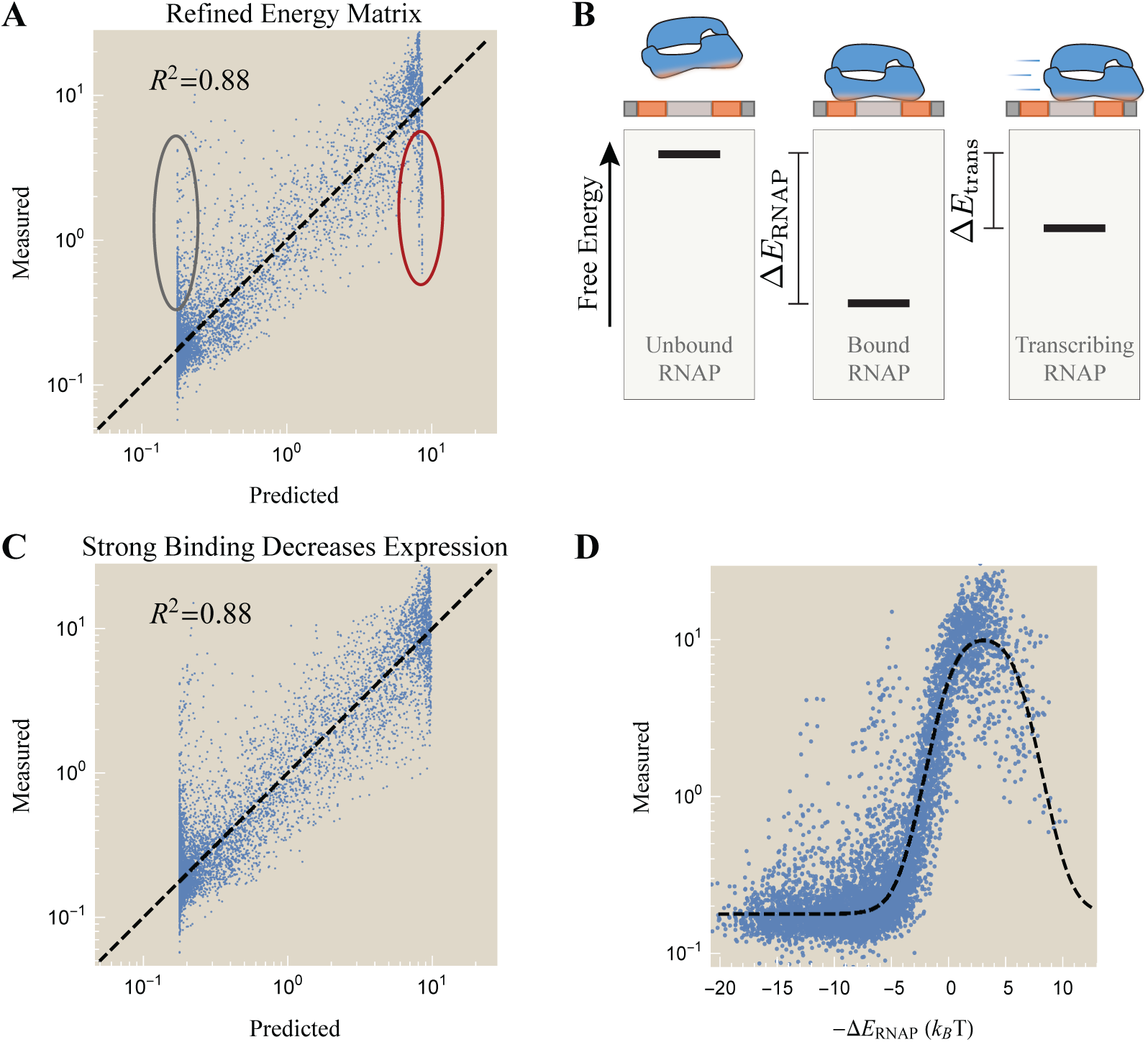
Gene expression is diminished for promoters that bind RNAP too tightly. (A) The refined energy matrix model (Eq. 5) characterizing all promoters, assuming a constant maximum level of gene expression *r*_max_. (B) The average level of transcription modeled as a two state system where the bound RNAP state (with free energy Δ*E*_RNAP_ relative to the unbound state) can enter a transcription initiation state with free energy Δ*E*_trans_. (C) Gene expression characterized using the modified maximum level of gene expression using Eq. S16 with Δ*E*_trans_ = *-*6.2 *k*_*B*_*T* . (D) Measured gene expression versus the promoter strength Δ*E*_RNAP_ (stronger promoters on the right because of the minus sign). The dashed line shows the prediction of the refined energy matrix model modified using Eq. S16.

## F. Dynamics of RNAP with Avidity

### F.1 Probability of the RNAP States at Equilibrium

In this section, we derive the probabilities of the four RNAP states shown in Fig. 1C in equilibrium. RNAP may be unbound (concentration *U*), singly bound at the -35 site (*B*_-35_), singly bound at the -10 site (*B*_-10_), or bound to both sites (*B*_-35,-10_). These concentrations must obey detailed balance,

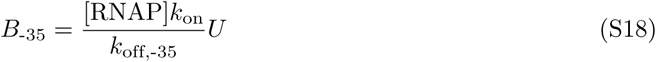

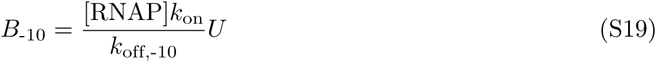

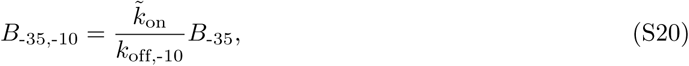

as well as the normalization condition

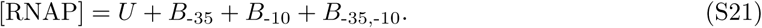

In writing Eqs. S18 and S19, we have assumed a sufficiently large reservoir of RNAP so that binding to the promoter of interest does not appreciably decrease the concentration of free RNAP (a reasonable assumption in *E. coli* where there are ≈ 2000 RNAP molecules (30)).

Eqs. S18-S21 can be solved to obtain the concentration of each RNAP state, namely,

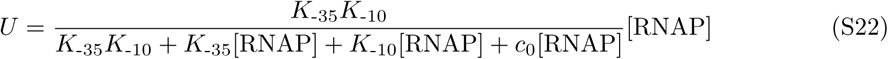

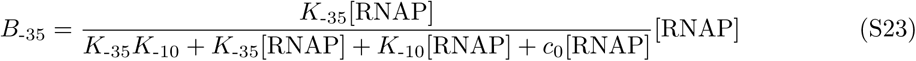

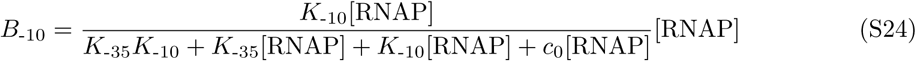

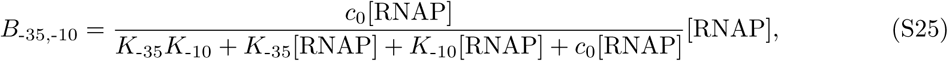

where we have defined the dissociation constants 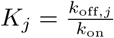 of free RNAP binding to the site *j* as well as the effective concentration 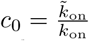 of singly bound RNAP binding to the remaining promoter site. If we further define the effective dissociation constant

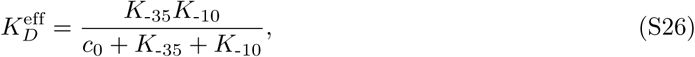

we can rewrite the probability of the unbound state as

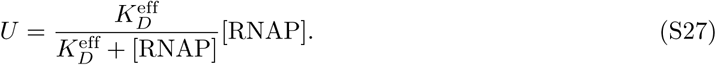

From this equation, we see that the promoter is bound 50% of the time 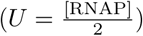 when 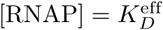, as stated in the main text.

### F.2 Dynamics of RNAP Unbinding with Avidity

Here, we rederive the results from the previous section by analyzing the dynamics of RNAP binding rather than its equilibrium configuration. This calculation highlights the intimate connection between the effective dissociation constant in Eq. S26 and the kinetics of RNAP binding.

**Figure S8.**
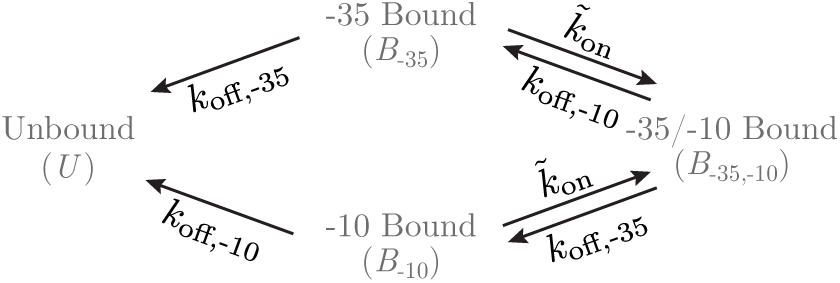
Dynamics of RNAP unbinding from the -35 and -10 sites. The avidity between the -35 and -10 sites will prolong the time before RNAP unbinds from the promoter.

To that end, we first compute the probability that a bound RNAP will remain bound after a time *t*. Since we are only interested in the unbinding process, we consider the rates diagram in Fig. S8 where the on-rates from the unbound state have been removed. Following Ref. (23), we assume that the three bound states – RNAP bound to only the -35 site (concentration *B*_-35_), only the -10 site (*B*_-10_), or to both sites (*B*_-35,-10_) – quickly equilibrate and compute the effective off-rate from these bound states to the RNAP unbound state (*U*). If the three bound states are in equilibrium, then there is no flux between any two states, namely,

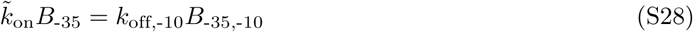

and

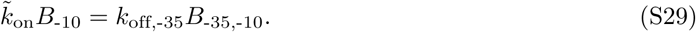

The total concentration of bound RNAP is given by

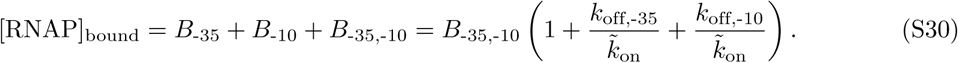

The loss of bound RNAP is caused by unbinding from the two singly bound forms, leading to the effective off-rate

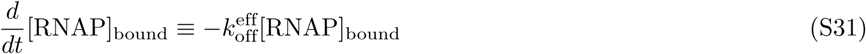

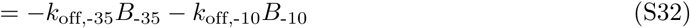

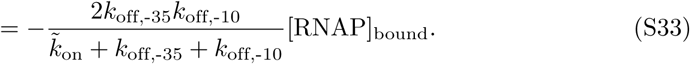

Hence, the dynamics of RNAP unbinding are characterized by

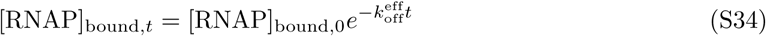

where the likelihood of remaining bound decreases exponentially according to the timescale 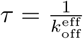.

Lastly, to connect this result to the calculations in the previous section, we return to the full model in Fig. 1C where unbound RNAP can associate onto the promoter. As in simple monovalent ligand-receptor systems, the effective dissociation constant Eq. S26 is related to the off-rate from the bound to unbound states 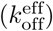 divided by the on-rate from the unbound to bound states (2*k*_on_), namely,

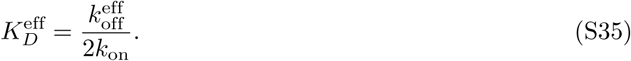

